# Most substitutions need more than one base change and an exact test tells which deposited scans measure the route

**DOI:** 10.64898/2026.11.17.767252

**Authors:** Taner Karagöl, Alper Karagöl

## Abstract

Design methods that reach a target through an intermediate take two substitutions at one position, and no deep mutational scan built from a single parental sequence reports the second. The detour is common, since 230 of the 380 ordered pairs of amino acids are more than one base change apart. Both substitutions are nevertheless present in the public record wherever one protein has been scanned on two parental backgrounds that differ at a position. We assemble every such comparison in MaveDB, 102,421 complete two-step routes over 65 protein clusters. The difference between a route and its two legs is a double difference, which is the epistasis between the tested position and the other positions at which the two backgrounds differ. Where the backgrounds differ nowhere else, that quantity is zero by construction. The rank correlation over the target residues must then return one, on any scale and against no null, which makes the comparison an exact test. Over the 741 qualifying pairs across 46 proteins the median is 0.965, and 34 of the 46 proteins return a median at or above 0.9. The arithmetic therefore holds across the population the record supplies, short natural domains drawn from families of near-identical sequences. Which pairs fail is not written in the record, since pairs drawn from one experiment set return 0.34 and pairs from unrelated submissions 0.97. The null such comparisons are read against fails the same test, returning +0.227 where it must return zero, more than the shift the comparison reports. Matching the control positions on the dispersion of the effects ranked halves that bias to +0.104. Two deposits of one protein agree at 0.642 where both score the same position. That bounds any predictor tested against one deposit at 0.80 for a position drawn at random, and at 0.90 and 0.98 where positions are scored more reliably. The test costs one correlation and settles, before any analysis, whether two deposits can be compared.

## Introduction

A substitution that a design method cannot reach in one step may be reachable in two. Converting a buried leucine to aspartate, or installing a histidine where the wild type carries a hydrophobe, can pass through an intermediate residue that is tolerated at the site, and the intermediate is often alanine, the residue that alanine-scanning mutagenesis has substituted at one position after another since the method was introduced [1]. The same route appears in the evolutionary record, where exchanges two base changes apart appear to proceed through a residue one change from each end [2,3], and where serine and threonine stand between the basic and acidic residues a direct exchange in the junctional adhesion molecules would have to cross [4]. Directed evolution reaches its target the same way, fixing the best variant found in each round of single-site mutagenesis and mutating again from there, so the walk is path dependent and its efficiency turns on the order in which the beneficial steps are found [5]. In silico design does the same across pairs of positions, pairing a substitution a model scores as deleterious with a compensatory one chosen in its presence [6]. Whether the route costs anything beyond the sum of its two steps is a quantitative question about the position, and answering it requires the effect of the second substitution measured from a protein that already carries the first.

There is every reason to expect that it might. The effect of a substitution depends on the residues around it, and that dependence is pervasive and not exceptional [7], and across the homologues of two structurally near-identical isoforms the residue variation at one position covaries with the variation at another [8]. Across the trajectories linking five substitutions in a beta-lactamase, most are closed to selection because individual mutations stop being beneficial in some combinations [9]. Over hundreds of millions of years of divergence in one domain family, the effects of most mutations become decorrelated from the effects they had at the start [10], and substitutions that were near neutral when they occurred grow steadily harder to revert [11]. In a viral protein whose variants are separated by many substitutions, the sites whose mutational costs have shifted are enriched near the positions at which the backgrounds differ [12]. A position’s response to a second substitution, after a first has been made there, is the smallest case of the same phenomenon, and it is the one case that a scan of single mutants cannot see.

Deep mutational scanning assays as many as a million variants of a protein in a single experiment [13], and the deposits now number in the millions of measurements [14], and neither supplies this one. A library is built from one parental sequence, and every variant in it carries the parental residue at every site it does not mutate. A variant whose parent at the site of interest is the intermediate is therefore absent, whatever the variant’s total substitution count, and the absence is structural and not an oversight of any one study. Libraries that carry many multi-substitution variants do not close the gap, because those variants place their substitutions at different positions. The gap matters because these measurements are the standard against which computational predictors of variant effect are scored [15], so a quantity absent from them is absent from the evidence those predictors are judged on. Where the two diverge has been mapped across forty human proteins, with the noise of the assay among the causes [16]. They also carry weight outside the laboratory, as the functional evidence a clinical variant classification may rest on [17], and several deposits of one gene are now combined to strengthen it [18], and the scores a deposit does not carry are imputed from models trained across domains [19]. Mutational effects on stability are largely conserved between homologous domains [20], while one enzyme compared in two orthologous backgrounds carries a minority of substitutions whose effect does not transfer [21], which makes whether two deposits can be put side by side a question with an answer instead of an assumption.

Combinatorial libraries over a fixed set of loci do not close it either. Such libraries are how empirical fitness landscapes have been built [22]: every combination of a small set of mutations, measured together. All single and double mutants of a protein domain give the first two steps of every pathway over its positions [23], a combinatorially complete library at three positions gives the effect of each substitution in hundreds of backgrounds [24], and four sites taken over all twenty residues show that paths blocked by sign epistasis can be circumvented by gaining and then losing a mutation [25]. Measuring several molecular phenotypes in several genetic backgrounds inside one experiment, and fitting a thermodynamic model to the result, maps the couplings between positions directly [26], which is the design the double difference asks for. A library of that last kind does carry the second step at a position, since every residue is present at every one of its sites, and the second leg is then the difference between two of its measured variants. That is available at the handful of sites the design fixes, in one protein, and it is a difference taken inside one experiment and not an independent measurement of the step. The largest such library for a viral protein assays all 2^15^ combinations of the fifteen receptor binding domain substitutions separating the Wuhan Hu-1 strain from Omicron BA.1 [27,28], which maps interaction between those positions exhaustively. Each position in that design takes two states, the Wuhan residue or the BA.1 residue, and reads carrying any third residue at a mutated position are discarded before analysis [27]. The question of what a position does after one substitution has been made there, and a different one is then attempted, falls outside it.

The measurement exists in the deposited record for a different reason. When one protein is scanned on two parental backgrounds that differ at a position, that position starts from a different residue in each scan. One scan gives the full spectrum of substitutions from the first residue, the other from the second, and if the change between the two parental residues is itself in the first spectrum, then the one-step route and the two-step route through the other background’s residue are both measured.

A second reason to expect a discrepancy is that the measured phenotype need not be linear in the property the substitutions act on. Where mutations are additive on an unobserved trait that is then passed through a monotonic nonlinear function, sums of measured effects will not match measured effects of combinations [7,29], and epistasis of that kind was detected in up to a third of multiple-mutation genotypes in a scan of one fluorescent protein [30]. A statistic that is unchanged by a monotonic rescaling separates this from a genuine failure of composition. The rank correlation used here is one, and where two deposits are related affinely and not merely monotonically, the product-moment correlation is admissible as well and is reported beside it.

Deposits of one protein that overlap in the variants they measure have been compared before, and ranking them against one another recovers a shared ordinal signal across assays of different scale and readout [31]. What such a comparison cannot supply is the value it ought to return, so a low correlation between two deposits is a fact about them and not a fault in either. The expansion supplies that value for part of the record. We assemble every such comparison in MaveDB [32] and ask whether substitution effects compose along the route. Expanding the three measurements answers that question before any data are read, for the pairs whose two backgrounds differ at the tested position and nowhere else, and what the expansion leaves is a test of the deposits rather than of the protein. We run it over every pair the record admits, report which proteins it rejects and what the deposited metadata fails to predict about them, show that the null these comparisons are ordinarily read against fails the same test by more than the effect it reports, and give the reliability the same control positions imply for anything scored against a single deposit.

## Results

### How often the genetic code forces a route through an intermediate

Consider one position of one protein, where a library built from a single parental sequence measures the effect of every target residue against the one residue that parent carries. If a method installs residue Z at that position and then replaces it with Y, the second of those two substitutions is made against Z, and no variant in that library carries Z as its parent at that site (Figure 1a). The library does determine a value for the step, as the difference between the effect of Y and the effect of Z, both measured against the parental residue X. That subtraction is the assumption under test here, and whether it holds is a question about the protein the intermediate is installed in rather than about the arithmetic.

**Figure 1.**
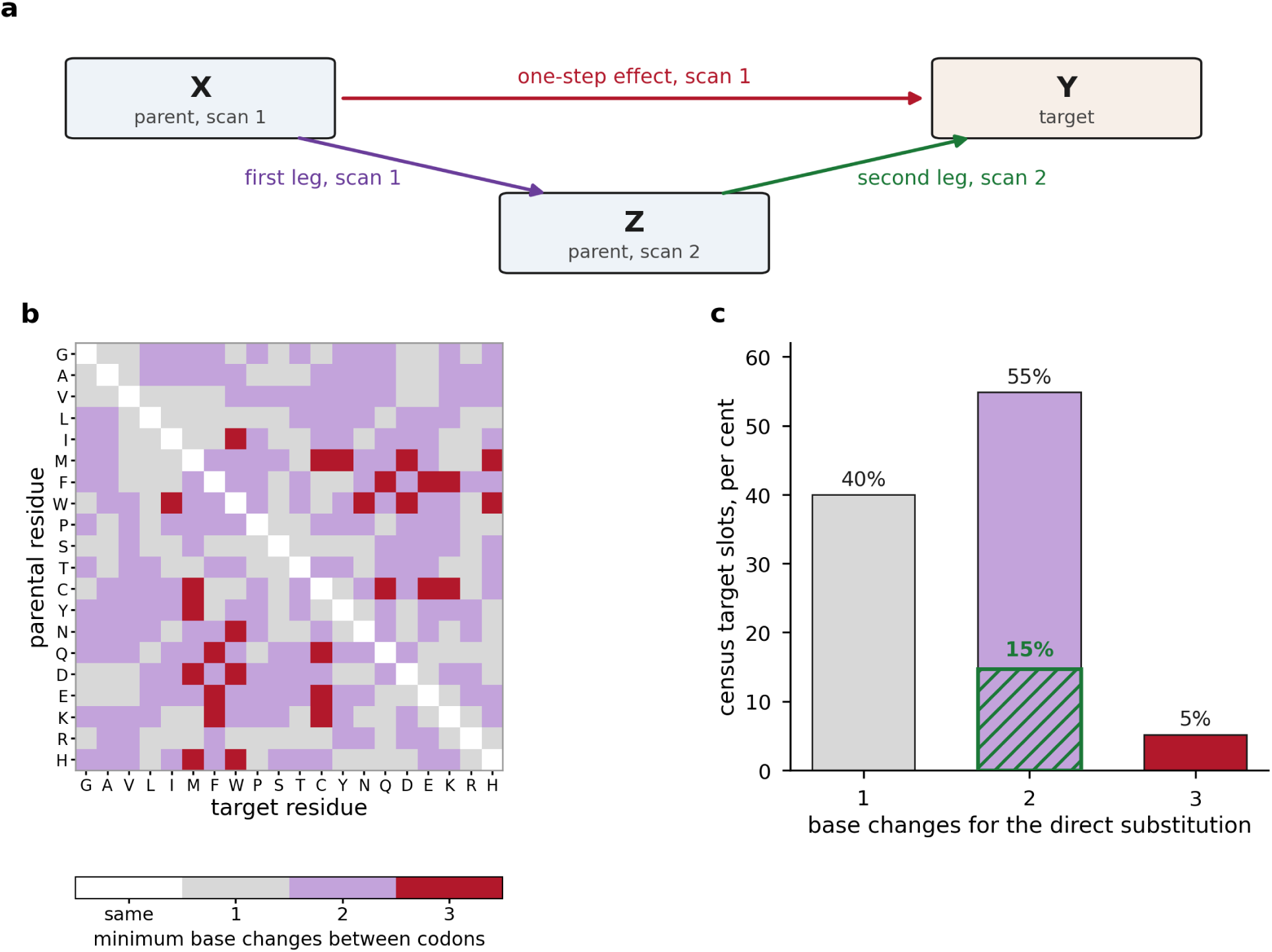
The route, what measures it, and how often the genetic code requires one. (a) At one position the first background carries X and the second carries Z. A library built on the first supplies the one-step effect of X to Y and the first leg of X to Z; the second leg of Z to Y is supplied only by a library whose parent at that position is already Z. Composition predicts that the one-step effect equals the sum of the two legs, and the residual is the difference. (b) Minimum number of base changes between the codons of every ordered pair of amino acids in the standard genetic code, minimised over synonymous codons, with residues ordered by side-chain character and not alphabetically. (c) Census target slots by the number of base changes the direct substitution requires. The hatched overlay, on the same axis, is the share of all slots whose direct substitution needs two base changes while each of the two legs needs one, so the route reaches the target by steps a single mutation can take and the direct substitution cannot.

The genetic code makes the detour common rather than exotic. A single nucleotide substitution carries the parent codon to the target codon only when the two differ at one base, and over the 380 ordered pairs of distinct amino acids in the standard code 150 pairs are one base change apart, 202 are two and 28 are three. Three pairs in five, 230 of 380, cannot be crossed by one mutation (Figure 1b), and the architecture of the code constrains which exchanges adaptation can explore [33]. The pattern is not uniform across the chemistry: exchanges within a size or polarity class are often one base change, and the exchanges a designer reaches for between classes are frequently not. The QTY and reverse-QTY conversions between leucine and glutamine, isoleucine and threonine, and phenylalanine and tyrosine are design codes built inside the reachable set, each a single base change at the second codon position [34,35]. A pair two base changes apart can still be coupled, and threonine and valine frequencies track one another in homologous sequences despite the double change that separates them [36].

### How often the detour is forced across the whole census

The same arithmetic over the census fixes how much of this resource the question touches. The census yields, once redundant deposits of one target are collapsed onto distinct sequences, 5,847 route fans, one for each sequence pair and position at which a complete route exists. They present 105,246 target slots once every target residue other than the parental and intermediate residues is counted. Of those, 42,065 sit one base change from the first background’s residue, 57,742 sit two and 5,439 sit three, so 63,181 of them, 60.0 per cent, are out of reach of a single mutation from the parent (Figure 1c). The first leg is itself a single base change at 4,108 of the 5,847 fans.

In 15,478 slots, 14.7 per cent of the total, the direct substitution needs two base changes while each leg of the route needs one. The route reaches a target that one mutation cannot while every step along it is one mutation, which is the configuration a stepwise process is confined to and a designed library is not. Counting every slot whose direct distance exceeds both of its legs gives 20,073, 19.1 per cent. Whether the two legs compose is therefore not a question about a modelling convenience. It decides whether the measured cost of the direct substitution is informative about the only path available to a process that moves one base at a time.

### The resource, and what defines a measurable route

As of 11 November 2026, MaveDB deposits 3,170 target gene records carrying 1,241 distinct target sequences. Two deposits form a background pair when their target sequences have the same length and differ at one or more positions by no more than 25.0 per cent of their residues, which is the condition under which positions correspond without an alignment. The census finds 586 such pairs, reads 688 score sets with 0 unreadable, and examines 49,562 position-and-direction combinations, of which 47,782 yield at least one route.

A route is complete when three measurements are present: the one-step effect of substituting the target residue Y for the first background’s residue X, the first leg from X to the second background’s residue Z, both in the first scan, and the second leg from Z to Y in the scan whose parent at that position is Z. Collapsing redundant deposits of one target onto distinct sequences gives 102,421 complete routes over 568 background pairs and 65 protein clusters, with 12,230 of them routed through alanine (Table 1). Each route is measured at one intermediate, the residue the other background happens to carry, so a result speaks to that intermediate and not to the best available one.

**Table 1.** The deposited resource and the routes it yields.

| **Quantity**<sup>1</sup> | **Value** |
|---|---|
| Target gene records read | 3,170 |
| Distinct target sequences | 1,241 |
| Background pairs at equal length and at most 25.0 per cent divergence | 586 |
| Score sets read | 688 (0 unreadable) |
| Position-and-direction combinations examined | 49,562 |
| Of those, yielding at least one route<sup>2</sup> | 47,782 |
| Complete routes, deduplicated onto distinct sequences<sup>3</sup> | 102,421 |
| Background pairs by sequence | 568 |
| Protein clusters contributing a route | 65 |
| Routes with alanine as the intermediate | 12,230 |

<sup>1</sup>Every value is produced by the census scripts and read from the derived tables; none is entered by hand.
<sup>2</sup>A combination yields a route when the first leg and at least one complete target are measured in both deposits.
<sup>3</sup>One deposited target sequence is often carried by several score sets, for instance a binding set and an expression set. Deduplication counts such a target once, so this value is smaller than the raw route count.

### How the routes are distributed across the record

The routes are not spread evenly over the clusters. The largest carries 32,471 of them, 31.7 per cent, the two largest together carry 58.6 per cent, and the median cluster carries 144 (Figure 2a). Any statement about the resource as a whole is therefore weighted towards a few heavily deposited proteins, which is why the tests are run within a protein system and combined across systems instead of pooled over routes.

**Figure 2.**
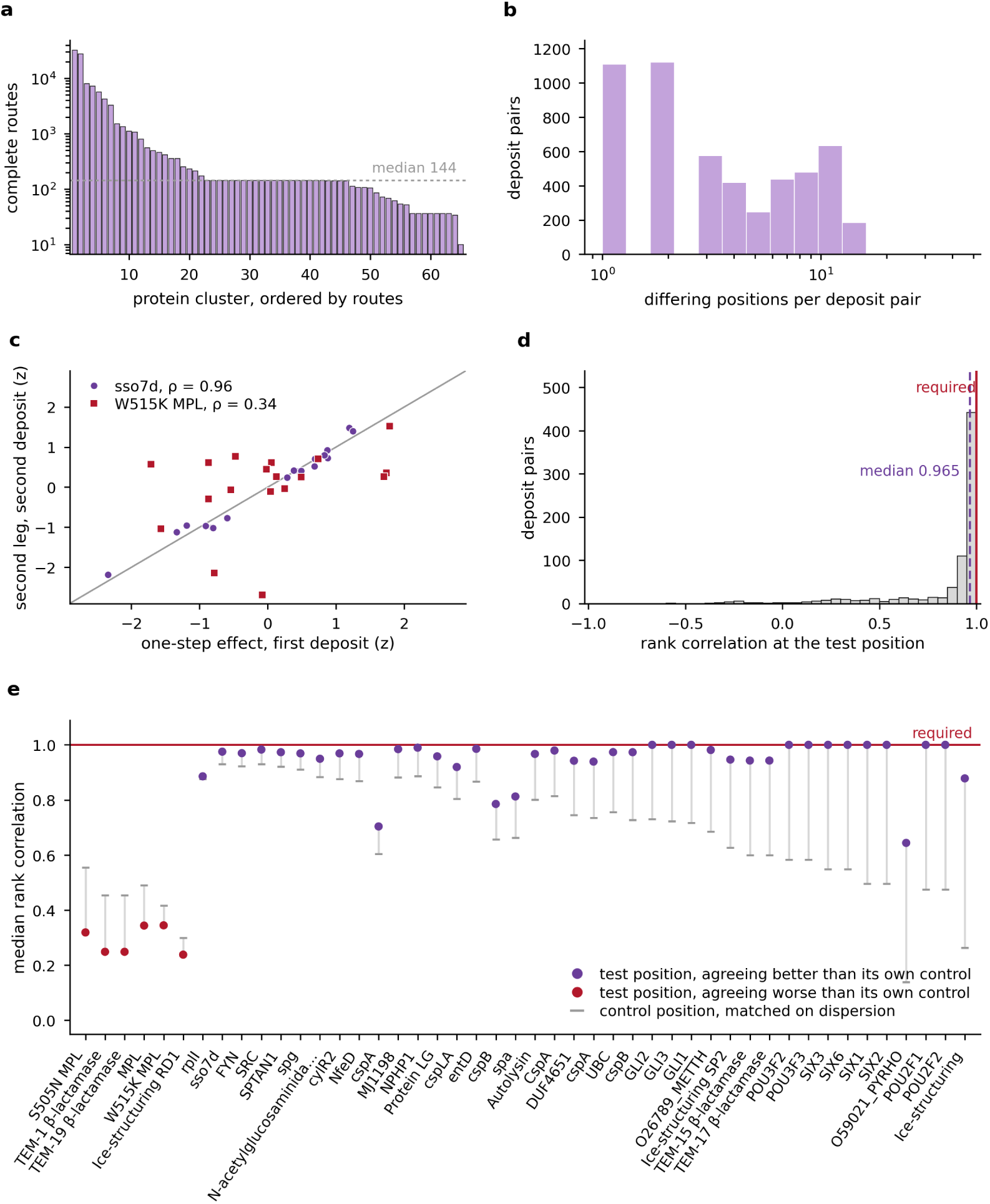
The deposited resource, and what the exact test returns across it. (a) Complete two-step routes by protein cluster, after collapsing redundant deposits of one target onto distinct sequences, ordered by size on a logarithmic axis. (b) Deposit pairs by the number of positions separating their two backgrounds. A pair differing at one position is the case in which the position under test is the only difference between the two proteins being compared. (c) Two deposit pairs whose target sequences differ at the position under test and nowhere else, at that position. The one-step effect from the first deposit and the second-leg effect from the second are standardised within each pair, since the test is invariant to any increasing rescaling of either, and the line is the relation the algebra requires. The sso7d pair sits on it; the W515K MPL pair, at the position its two backgrounds differ at, does not. (d) The rank correlation at the test position over all 741 such pairs, against the value of one the algebra requires of every one of them. (e) One protein at a time, the median rank correlation at its test positions and at its control positions, where the parental residue is the same in both deposits and which are matched to the test position on the dispersion of their effects. Both are required to be one. Proteins are ordered by the difference between them, as Table 2 is, so the 6 that agree worse where the route is assembled stand at the left and are marked.

**Table 2.** The exact test, one protein at a time.

| **Protein**<sup>1</sup> | **Pairs** | **Median** *ρ*<sup>2</sup> | **Median r**<sup>3</sup> | **Above 0.9** | **Paired difference**<sup>4</sup> | **p**<sup>5</sup> |
|---|---|---|---|---|---|---|
| S505N MPL | 23 | 0.319 | 0.356 | 1 | −0.235 | 1.4e−7 |
| TEM-1 *β*-lactamase | 11 | 0.248 | 0.151 | 0 | −0.206 | 1.0e−5 |
| TEM-19 *β*-lactamase | 11 | 0.248 | 0.151 | 0 | −0.206 | 1.0e−5 |
| MPL | 45 | 0.344 | 0.338 | 1 | −0.146 | 5.4e−10 |
| W515K MPL | 22 | 0.345 | 0.320 | 0 | −0.071 | 8.0e−4 |
| Ice-structuring protein RD1 | 9 | 0.238 | 0.992 | 0 | −0.062 | 1.2e−2 |
| rplI | 27 | 0.886 | 0.956 | 10 | +0.010 | 3.5e−7 |
| sso7d | 9 | 0.975 | 0.995 | 9 | +0.045 | 2.6e−2 |
| FYN SH3 domain | 36 | 0.970 | 0.993 | 36 | +0.047 | 5.7e−6 |
| SRC | 18 | 0.983 | 0.989 | 14 | +0.052 | 1.1e−2 |
| SPTAN1 | 72 | 0.973 | 0.993 | 70 | +0.053 | 2.7e−7 |
| spg | 99 | 0.969 | 0.983 | 90 | +0.059 | 9.9e−2 |
| N-acetylglucosaminidase | 18 | 0.949 | 0.980 | 14 | +0.067 | 1.8e−1 |
| cylR2 | 18 | 0.969 | 0.991 | 18 | +0.093 | 3.9e−1 |
| NfeD domain-containing protein | 18 | 0.967 | 0.983 | 17 | +0.098 | 3.8e−1 |
| cspA Cold-shock protein, DNA-bi… | 6 | 0.704 | 0.751 | 3 | +0.099 | 6.7e−1 |
| Uncharacterized protein MJ1198 | 18 | 0.985 | 0.990 | 18 | +0.102 | 8.9e−1 |
| NPHP1 | 18 | 0.990 | 0.993 | 18 | +0.104 | 9.1e−1 |
| Protein LG | 18 | 0.958 | 0.986 | 13 | +0.113 | 5.6e−1 |
| cspLA | 18 | 0.919 | 0.947 | 9 | +0.116 | 2.2e−1 |
| entD | 36 | 0.986 | 0.990 | 32 | +0.119 | 5.6e−1 |
| cspB Cold-shock protein, DNA-bi… | 6 | 0.785 | 0.847 | 3 | +0.128 | 8.0e−1 |
| spa | 18 | 0.813 | 0.924 | 5 | +0.150 | 3.0e−1 |
| Autolysin | 18 | 0.967 | 0.987 | 17 | +0.166 | 1.6e−2 |
| Cold shock-like protein CspA | 18 | 0.979 | 0.985 | 18 | +0.166 | 1.8e−3 |
| DUF4651 domain-containing prote… | 9 | 0.942 | 0.980 | 9 | +0.197 | 1.3e−1 |
| cspA | 24 | 0.939 | 0.981 | 18 | +0.205 | 9.8e−3 |
| UBC | 72 | 0.974 | 0.931 | 53 | +0.218 | 5.0e−9 |
| cspB | 24 | 0.973 | 0.989 | 21 | +0.247 | 1.6e−4 |
| GLI2 Zinc finger C2H2-type doma… | 1 | 1.000 | 1.000 | 1 | +0.269 | n/a |
| GLI3 Zinc finger C2H2-type doma… | 2 | 1.000 | 1.000 | 2 | +0.277 | n/a |
| GLI1 Zinc finger C2H2-type doma… | 1 | 1.000 | 1.000 | 1 | +0.284 | n/a |
| O26789_METTH | 18 | 0.981 | 0.997 | 18 | +0.296 | 4.2e−7 |
| Ice-structuring protein SP2 | 18 | 0.946 | 0.973 | 15 | +0.319 | 9.1e−7 |
| TEM-15 *β*-lactamase | 1 | 0.943 | 0.958 | 1 | +0.343 | n/a |
| TEM-17 *β*-lactamase | 1 | 0.943 | 0.958 | 1 | +0.343 | n/a |
| POU3F2 Homeobox domain | 1 | 1.000 | 1.000 | 1 | +0.417 | n/a |
| POU3F3 Homeobox domain | 1 | 1.000 | 1.000 | 1 | +0.417 | n/a |
| SIX3 Homeobox domain | 1 | 1.000 | 1.000 | 1 | +0.452 | n/a |
| SIX6 Homeobox domain | 1 | 1.000 | 1.000 | 1 | +0.452 | n/a |
| SIX1 Homeobox domain | 1 | 1.000 | 1.000 | 1 | +0.504 | n/a |
| SIX2 Homeobox domain | 1 | 1.000 | 1.000 | 1 | +0.504 | n/a |
| O59021_PYRHO | 18 | 0.644 | 0.873 | 0 | +0.505 | 1.2e−10 |
| POU2F1 POU-specific domain | 1 | 1.000 | 1.000 | 1 | +0.525 | n/a |
| POU2F2 POU-specific domain | 1 | 1.000 | 1.000 | 1 | +0.525 | n/a |
| Ice-structuring protein | 9 | 0.878 | 0.961 | 2 | +0.615 | 1.5e−6 |

<sup>1</sup>Every protein carrying at least one deposit pair whose two target sequences differ at the position under test and nowhere else. Rows are ordered by the paired difference, so the proteins that agree worse where the route is assembled come first.
<sup>2</sup>The Spearman rank correlation between the one-step vector and the second-leg vector over the target residues, which the algebra fixes at 1 for every row of this table.
<sup>3</sup>The Pearson correlation over the same vectors, which the same algebra also fixes at 1, since each deposit reports an increasing affine function of the underlying effect.
<sup>4</sup>The rank correlation at the test position minus the rank correlation the same deposit pair reaches at control positions matched to it on the dispersion of their effects, where the value is also 1. A negative difference is a failure specific to the assembled route rather than a deposit pair that is noisy throughout.
<sup>5</sup>Two-sided rank-sum p for this protein's paired differences against those of every other protein, which is the comparison the question asks for, since the difference is not expected to be zero. Pairs within a protein share deposits, so the value is optimistic. A protein carrying fewer than three pairs is marked n/a, since a rank-sum test over so few is not informative.

How many deposits a background pair draws on sets how much data a protein system contributes, and how many positions separate its two backgrounds sets how much of the protein differs between the scan that supplies the one-step effect and the scan that supplies the second leg. A pair differing at one position alone is the cleanest case available, because the position under test is then the only difference between the two proteins being compared (Figure 2b).

### The discrepancy is a double difference, and one case fixes its value

Expanding the three measurements shows what the routing residual contains. Write the phenotype of a sequence as a function of the residue at the tested position and of the rest of the protein. The first background is (X, B) and the second is (Z, B^′^), and the residual at target Y is

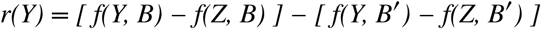

The routing residual is a double difference: the effect of substituting Y for Z measured on one background, minus the same effect measured on the other. It is therefore the epistasis between the substitution at the tested position and the positions at which the two backgrounds differ elsewhere, and it is nothing besides that. The first leg and the parental residue X both cancel. Taken over all substitutions instead of at one position, the same difference between two scans of homologous backgrounds is what a joint model of several deposits infers as a shift in mutational effect [37].

### The two cases whose value the expansion fixes

When the two backgrounds differ at the tested position and nowhere else, B and B^′^ are the same protein, the two brackets are the same quantity, and r(Y) is zero at every target. The second leg is then the one-step effect minus a constant, so in a common unit the two vectors lie on a line of unit slope. Each deposit reports its own increasing rescaling of that unit, which moves the slope and the intercept and leaves the ordering alone, so what survives for deposits of unknown scale is that their rank correlation is one. It is an identity about the protein, and what it leaves is a statement about the measurement.

A second case follows from the same expansion and costs nothing extra. At a position whose parental residue is the same in both deposits, no routing arises at all: the two scans measure one set of substitutions twice, against one parent, and their rank correlation must be one as well. The first case is a route assembled across two deposits; the second is the same substitution measured twice. The value fixed in advance is one in both, at the tested position and at a position whose parental residue the two deposits share (Figure 2c).

### An exact test for a route assembled across two deposits

The first case is a test, not an observation. Any pair of deposits whose target sequences differ at the position under test and nowhere else carries a quantity whose value is known in advance, so a correlation between the one-step vector and the second-leg vector over the target residues can be computed and compared with one. It needs no null, no replicate, no calibration between the two deposits and no assumption about the scale either reports, and it costs one correlation per position. Nothing about the protein is assumed either: the identity holds whatever the epistasis at that site, because the background that would carry it is the same protein on both sides.

Applying it to the whole census finds 796 deposit pairs of the 4,831 that carry a complete route whose two translated target sequences differ at exactly one position, and whose differing position is the one under test. A route has a direction and the census counts routes that way, while the statistic is symmetric in the two deposits, so every count of comparisons from here on is of unordered pairs. Reading the 384 score sets behind them and requiring at least 6 target residues measured in both leaves 741 pairs across 46 proteins, every one of them an assembled route whose answer is fixed at one. Most of them are short natural domains whose deposits report folding stability measured by cDNA display proteolysis [38], and the census finds them in families of near-identical sequences, which is what supplies a pair differing at one position. The deposits carry no publication identifier, so the assay has to be read off the method text they do carry. Equality outside the tested position is checked on the deposited target sequence, which is the protein the assay reports on rather than always the whole protein assayed, so two deposits agreeing across a scanned window may differ outside it. The admitted targets are short, a median of 62 residues with 729 of 796 at 100 or fewer, and the record identifies both deposits of only 61 of those pairs by accession at all.

### Which correlation the expansion fixes, and what passing establishes

Which correlation to use is settled by the same expansion. Each deposit reports an increasing affine function of the underlying effect, so the two reported vectors are affinely related and the product-moment correlation is fixed at one as well, not only the rank correlation. The product-moment form tests the affine prediction itself and is the more powerful of the two where that prediction holds; the rank form tests only the ordering and survives a monotone but nonlinear difference between the deposits, such as one readout saturating where the other does not. The rank correlation is reported throughout as the conservative choice, and the product-moment and Kendall correlations agree with it (Supplementary Table S9). The test is necessary and not sufficient, since a rank correlation is unmoved by any increasing map, so a pair that returns one has shown only that the two deposits order the substitutions alike at that position, and the product-moment form shows the relation to be affine while leaving its slope and intercept free, so neither fixes the two deposits in a common unit. The statistic is blind by construction to a penalty that falls equally on every target, so a position whose route carries a uniform cost is reported here as composing.

Most of them are consistent, at a median rank correlation of 0.965, a lower quartile of 0.897, and 552 of 741 pairs exceeding 0.9 (Figure 2d, Supplementary Table S13). The product-moment correlation over the same pairs has a median of 0.984 with 607 above 0.9, so the affine prediction is met at least as well as the weaker one: 69 pairs clear 0.9 on the product-moment correlation while falling below it on the rank correlation, against 14 the other way. Taken one protein at a time, 34 of the 46 return a median at or above 0.9 and 12 fall below it (Table 2, Figure 2e). The arithmetic that assembles a two-step route out of two separate deposits therefore reconstructs a consistent ordering for most of the deposited record, and the exceptions are specific, not general. That median describes this population, short natural domains in families of near-identical sequences, more closely than it describes MaveDB as a whole.

### What the test rejects, and why an absolute threshold is wrong

An absolute threshold is the wrong criterion, because the same two deposits must also return one at every position whose parental residue they share, and most pairs fall short there too. What isolates a failure specific to the assembled route is the paired comparison: the correlation at the test position minus the correlation the same deposit pair reaches at control positions matched to it on the dispersion of their effects. That difference has a median of +0.108 across the 741 pairs, so a test position is ordinarily measured better than the controls standing in for it, and a protein whose difference is negative is agreeing worse exactly where the route is assembled. 6 of the 46 proteins are negative, and every one of them departs from the rest of the census on a rank-sum test of its paired differences, at a Benjamini-Hochberg threshold of 2.59e−2 across the 46 proteins (Table 2). Pairs within a protein share deposits, so those p values are optimistic.

Those proteins are not a random handful, since three of them are the thrombopoietin receptor on its wildtype, S505N and W515K backgrounds, at medians of 0.34, 0.32 and 0.34 over 90 pairs, at paired differences of −0.15, −0.24 and −0.07. Their deposits come from 2 experiment sets, of which 1 carries a publication identifier, covering the wild-type and S505N screens [39]. The set holding the W515K background and the drug-dose screens is deposited without one. Two more are TEM-1 and TEM-19 *β*-lactamase, both at 0.25, at −0.21 and −0.21, the two largest negative differences in the census, whose deposits list a series of studies along one adaptive path [40,41,42,43,44]. They are also the two a reader would reach for first, because each is a protein scanned on backgrounds that a point substitution defines, which is the configuration this measurement calls for. A system here is a set of deposit pairs sharing a base target name, which groups engineered variants of one protein with its wild type and is not a clade, a family or a fold, so these three entries are one receptor on three backgrounds.

At one differing position the routing residual is identically zero, so a departure measured there cannot be epistasis between the tested position and the rest of the protein, because the two backgrounds hold that rest identical. Whatever these systems are departing from, it is not the quantity the comparison is meant to measure, and a statistic that cannot separate the two will report the disagreement as a result.

### What the deposited record does not tell

The deposited identifiers do not predict which pairs fail (Supplementary Table S10). Pairs whose two deposits are drawn from one MaveDB experiment set return a median of 0.34 over 46 pairs, and pairs whose deposits are drawn from different experiment sets return 0.97 over 695. The relation runs the wrong way, because the deposits that the record groups together agree least. Exactly 1 of the 796 pairs has both deposits inside a single experiment, which is the one relation that guarantees a common assay by curation, and it fails at 0.35.

Nor does a mismatch in what is measured account for it. Classifying each pair as one assay where both deposits sit in one experiment or carry the same score set title once the target’s own name is removed, and as different assays otherwise, splits the 741 pairs into 223 and 518. Their medians are 0.973 and 0.957, close enough to each other that the split explains nothing, and the thrombopoietin receptor pairs that do sit inside one experiment fail as badly as the rest. What those deposits measure is a receptor made constitutively active, selected from a wild-type background in one and from an already-active S505N background in the other, so the range over which either screen can separate variants is not the same. A difference in where a readout saturates is not an increasing map, and it is the one kind of difference a rank statistic cannot absorb.

The practical consequence is that the record cannot be filtered on provenance. Two deposits grouped under one experiment set may be incomparable and two deposits from unrelated submissions may agree almost exactly, and nothing short of the test distinguishes them. The test is available for every pair the record contains, before any analysis is run on it. It applies, though, only where the two target sequences differ at the tested position and nowhere else, which is 796 of the 4,831 pairs carrying a complete route. For the remainder no value is known in advance, so what the majority of the resource supports is an open question.

### The null the comparison is measured against fails the same test

Nothing above needs a null, which is what makes it useful as a check on everything that does. A routing discrepancy is ordinarily read against the agreement the same two deposits reach at positions whose parental residue they share, matched to the test position on the number of target residues measured. Both quantities must return one at a pair that differs at a single position, so the shift between them must return exactly zero, and the construction can be audited instead of assumed, in the way a conservation score’s two failure modes can be computed from the alignment before the score is relied on [45].

It returns +0.227. Over the 741 pairs the median shift is that, the median absolute shift is 0.255, and the sign is consistent: the test position agrees better than its null in 665 of 741 pairs (Figure 3a). The null sits at a median of 0.658 where it too must return one.

**Figure 3.**
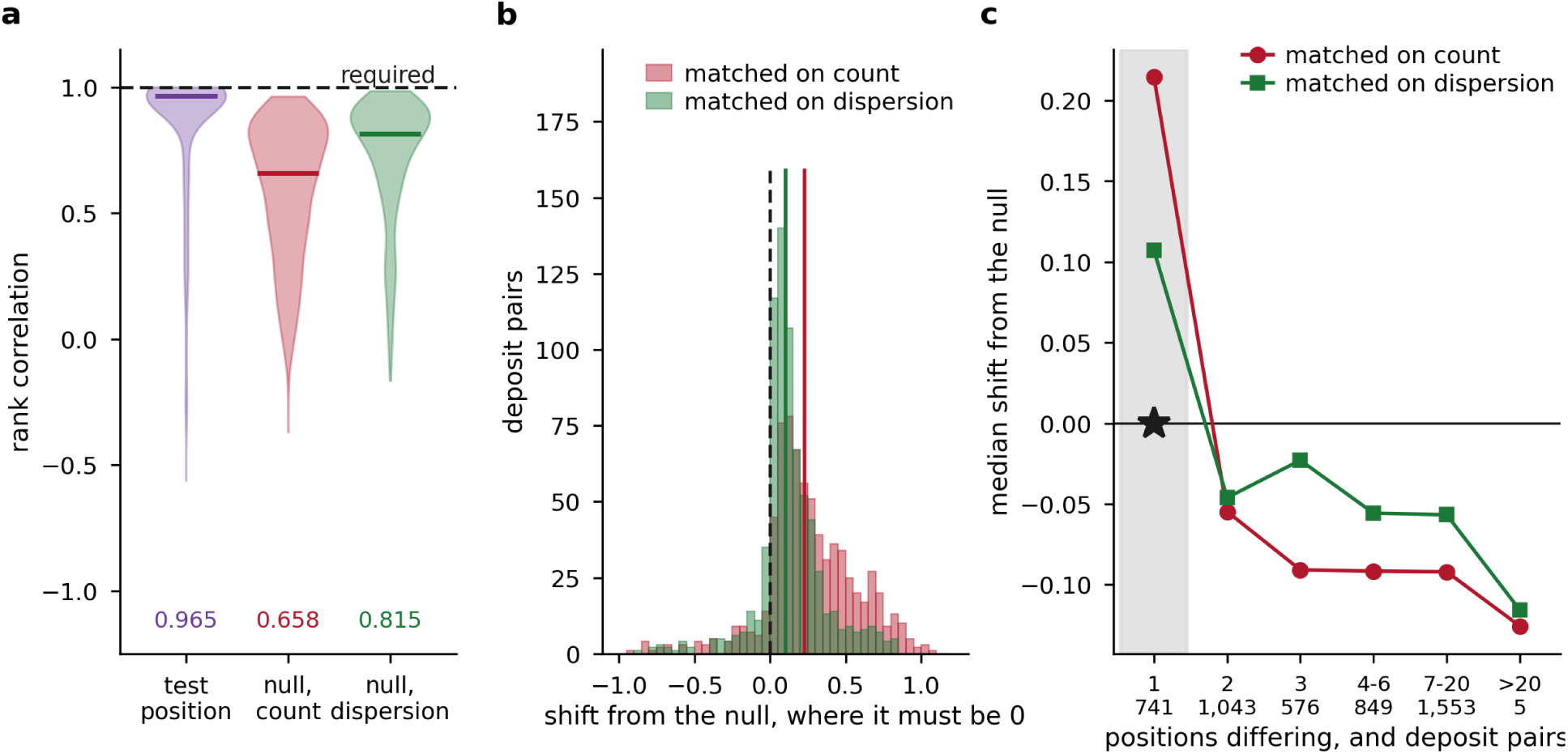
The null the same comparison is read against, measured where its value is known. (a) The rank correlation at the test position and at the control positions the null draws from, over the 741 pairs at one differing position, where all three must return one. The null matched only on the number of target residues sits furthest below it; matching the control positions on the dispersion of the effects being ranked recovers much of the gap. Bars are medians. (b) The shift between the test position and its null for the same pairs, which the algebra fixes at exactly zero. Vertical lines are medians. (c) The median shift against the number of positions at which the two backgrounds differ, over 4,767 deposit pairs. The shaded bin is the one whose value is known, where the star marks the zero the algebra fixes, and the trend across the remaining bins is flat.

### Where the bias comes from, and how much of it matching removes

The sign of the shift points to the cause, since a position at which two backgrounds differ is a position somebody chose to vary, and the substitutions there spread over a wider range of effect than at a position drawn at random from the same protein. A rank correlation between two noisy measurements rises with that spread, so a test position is measured more reliably than the control positions standing in for it, and the shift inherits the difference. Matching the null on the dispersion of the effects being ranked, as well as on their number, is enough to test it: drawing the control positions nearest in log standard deviation to the test position brings the dispersion ratio to a median of 0.95, lifts the null to 0.815 and halves the shift to +0.104, with the median absolute shift falling from 0.255 to 0.126 (Figure 3b).

Half the bias is therefore a matching failure with a known remedy, and half is not. What matters for anything built on this comparison is the size: a null that is wrong by 0.23 where the answer is zero cannot support a conclusion drawn from a shift smaller than that. What remains after matching is not identified, and part may be imperfect matching and part a second mechanism.

### The reported trend is the bias, measured where the answer is known

The prediction the double difference makes is graded, because the routing residual is the epistasis between the tested position and the positions at which the backgrounds differ elsewhere, so it should vanish when there are none and grow as there are more of them. Measuring that needs the whole census and not a summary over protein systems, and the shift can be computed for every deposit pair the census contains, binned on the number of positions separating the two target sequences.

Over 4,767 deposit pairs across 102 proteins the shift under the standard null runs +0.215 at one differing position, −0.055 at two, −0.091 at three, −0.092 from four to six and −0.092 from seven to twenty (Figure 3c). Correlated against the logarithm of the separation it gives −0.350, and −0.365 with the number of test positions held constant.

### The trend is carried by the one bin whose value is known

The trend is one bin, since from two separating positions to more than twenty the shift neither grows nor orders itself, staying between −0.13 and −0.05 across 4,026 pairs, and the whole correlation is carried by the first bin, which is the one bin whose value is fixed at zero by the algebra and returns +0.215 instead. A statistic that is wrong by that much where it can be checked, and flat everywhere it cannot, is not measuring a graded cost of background separation.

Removing that bin removes the trend, and over the 4,026 pairs whose backgrounds differ at two positions or more, where no value is known in advance, the correlation against the logarithm of the separation is −0.036, and +0.006 with the number of test positions held constant, so the claim does not rest on the binning (Supplementary Table S7).

### What matching the null leaves, and what the record lacks

Matching the null on dispersion does not rescue it. The same bins run +0.107, −0.046, −0.023, −0.056 and −0.057, the trend falls to −0.274, and the first bin carries it after the correction as well. The correction shrinks the artefact and leaves nothing graded behind. These deposits do not show the routing cost rising with the distance between the backgrounds, and the appearance of a trend comes from comparing test positions against control positions that are not their equals. Without a position whose answer is fixed in advance, the same gradient reads as a measured cost of separation, and nothing in the data separates the two. What this bounds is a measurement and not a demonstration that routes compose, because the shift carries a bias of +0.227 where its value is known, so an increase smaller than that would not be visible here, and whether routing costs grow with separation remains untested by these deposits.

Deposits with the structure this measurement needs do exist, and there are very few of them. The *β*-lactamase sets come from a series in which successive backgrounds along one adaptive path differ by single substitutions [40,41,42,43,44], and the thrombopoietin receptor sets scan one receptor on backgrounds that single clinical substitutions define [39]. Both were built for other purposes, both are what a direct test of routing requires, and both are among the 6 proteins the paired difference rejects. What the field lacks is not analysis of the deposited record but one scan of one protein on two backgrounds separated by one substitution, run as a single experiment on a single assay, which the test given here would then certify in a line.

### How far two deposits of one protein agree, and what that bounds

The control positions are a result in their own right and not only a null. At a position whose parental residue both deposits share, both scans score the same substitutions against the same parent, so their rank correlation must also be one, and no routing, no epistasis and no scale assumption enters. Across the same 741 pairs the median is 0.642, with a lower quartile of 0.444 and 45 pairs above 0.9 (Figure 3a). Two deposits of one protein order the substitutions at a position they both measure, against a parent they both carry, at about that much agreement.

That figure is a reliability, and it bounds what any method can score against these data. If two deposits are parallel measurements of one quantity with independent error, the correlation between either of them and the quantity itself is at most the square root of the correlation between them, which puts the ceiling for a predictor scored against a single deposit at 0.80 at a position taken at random. The relation is classical and is stated here on the rank scale as an approximation, since the exact result is a product-moment one and holds when the two errors are uncorrelated [46]. Two scans sharing a protocol or a reagent batch would have correlated errors, which inflates the agreement and raises the apparent ceiling. A reliability estimated instead from the replicate scores and standard errors a deposit carries bounds predictor benchmarks the same way, and most of MaveDB carries what such an estimate needs [47].

The ceiling moves with the positions scored, which is the part worth carrying away. At control positions matched to the test positions on the dispersion of their effects the agreement is 0.815 and the ceiling 0.90; at the test positions themselves it is 0.965 and 0.98. A benchmark assembled from well-measured positions and one assembled from an unselected sweep of a protein are not scored against the same ceiling, and a reported correlation that is not accompanied by one cannot be read as a distance from the truth. The positions the test examines are those at which somebody chose to make a second background, which is not a random sample. They are residues separating natural homologues in the proteolysis panels, positions that changed in a virus under immune pressure in the receptor binding domain, and positions a designer selected in the engineered systems. The control positions are the rest of each protein and closer to a sweep, which is why the two figures differ. A score read as a property of a position behaves the same way, and the two failure modes of alignment-based conservation, one inflating and one concealing constraint, are computable from the alignment before the score is used [45], and a conservation score conflates a residue’s structural rigidity with its functional motion until the two are separated [48]. The arithmetic above assumes the two deposits measure one quantity, which is exactly what the exact test establishes, and which is why it has to come first.

### A worked case with an independently verified scale

The SARS-CoV-2 receptor binding domain has been scanned on the Wuhan Hu-1 background [49] and on the Omicron BA.1 and BA.2 backgrounds [50], each for ACE2 binding and for surface expression. Scans of the later Omicron descendants, compared against these, find positions whose preferred residues are still changing from one background to the next [51]. Translating the three deposited target sequences and comparing them residue by residue gives 15 differing positions between Wuhan and BA.1, 16 between Wuhan and BA.2 and 6 between BA.1 and BA.2, for 18 distinct positions. The offset between the numbering of the deposits and spike numbering is solved for, not assumed: exactly one shift carries the Wuhan to BA.1 differences onto the recognised spike substitution names, and it is 330.

These six deposits are the only ones in the census whose scales were checked directly (Supplementary Table S4). The two Omicron sets deposit an absolute phenotype alongside the effect, and the effect equals that absolute value minus one parental constant for all but a single row per set, with parental values of 9.22224 and 9.25549 for binding and 9.73227 and 9.85355 for expression. All six are effects in the same units, which is what makes a sum of two of them dimensionally legitimate.

Of the 74 position-and-phenotype combinations, 15 depart from composition in the magnitude of the routing residual at a Benjamini-Hochberg adjusted p below 0.05, at spike positions 373, 376, 408, 477, 493, 496, 498, 501, 505 (Figure 5a, Supplementary Table S1). The result is unchanged across the whole censoring sweep, which costs at most 11 of 666 binding routes and none of the expression routes.

### One measured route, at spike 498

At spike 498 the first background carries glutamine and the second arginine, and the route to serine carries the largest routing residual of the 666 binding routes in the set. Substituting serine for glutamine costs 0.17 in log_10_ K_a_ in the Wuhan scan, installing the arginine costs 0.06, and substituting serine for that arginine in the BA.1 scan costs 2.96. The route is therefore +2.8493 worse than the direct substitution, against a null median of 0.323. Installing the arginine is nearly free and makes the position load-bearing, so the step away from it is close to disqualifying while the direct substitution is cheap. Entrenchment of this kind at spike 498 and 501, and its absence at 493, is reported in the study that produced the Omicron deposits [50].

This route is also an instance of the configuration counted above. Glutamine and serine are two base changes apart, while glutamine to arginine and arginine to serine are each one, so the detour is the only version of this substitution a process moving one base at a time can make, and it is the version the direct measurement misprices by +2.8493 (Figure 4).

**Figure 4.**
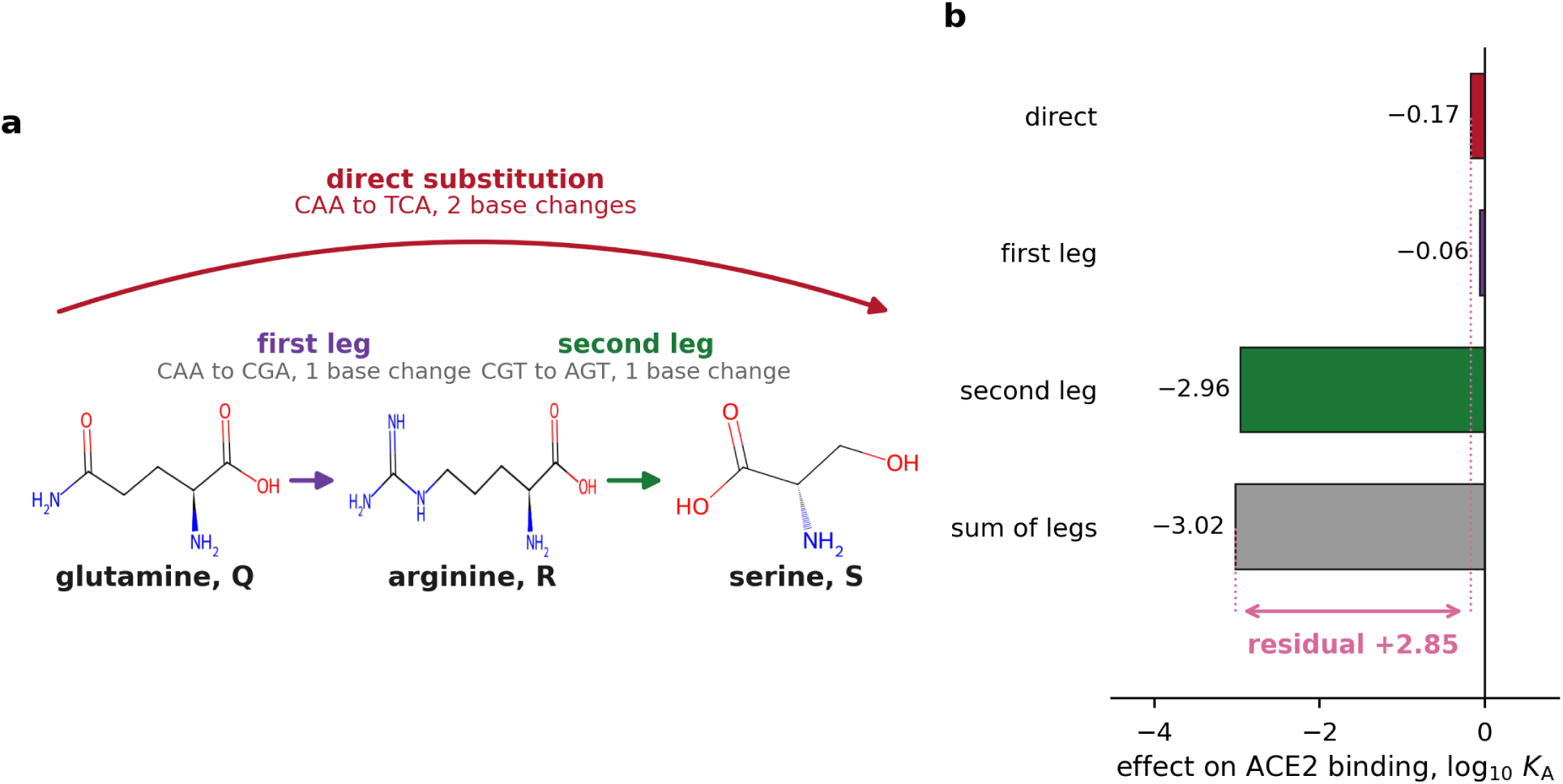
The chemistry of one measured route, at spike 498. (a) The first background carries glutamine at this position and the second carries arginine. Each step is drawn with the codon path that realises it. Glutamine to serine is two base changes and is measured directly in the Wuhan scan, while glutamine to arginine and arginine to serine are one base change each and are measured in different scans. (b) The direct effect, the two legs and their sum, with the routing residual spanning the gap between the direct effect and the sum of the legs. This is the largest residual of the binding routes in the set, and the position is one at which entrenchment was reported independently.

**Figure 5.**
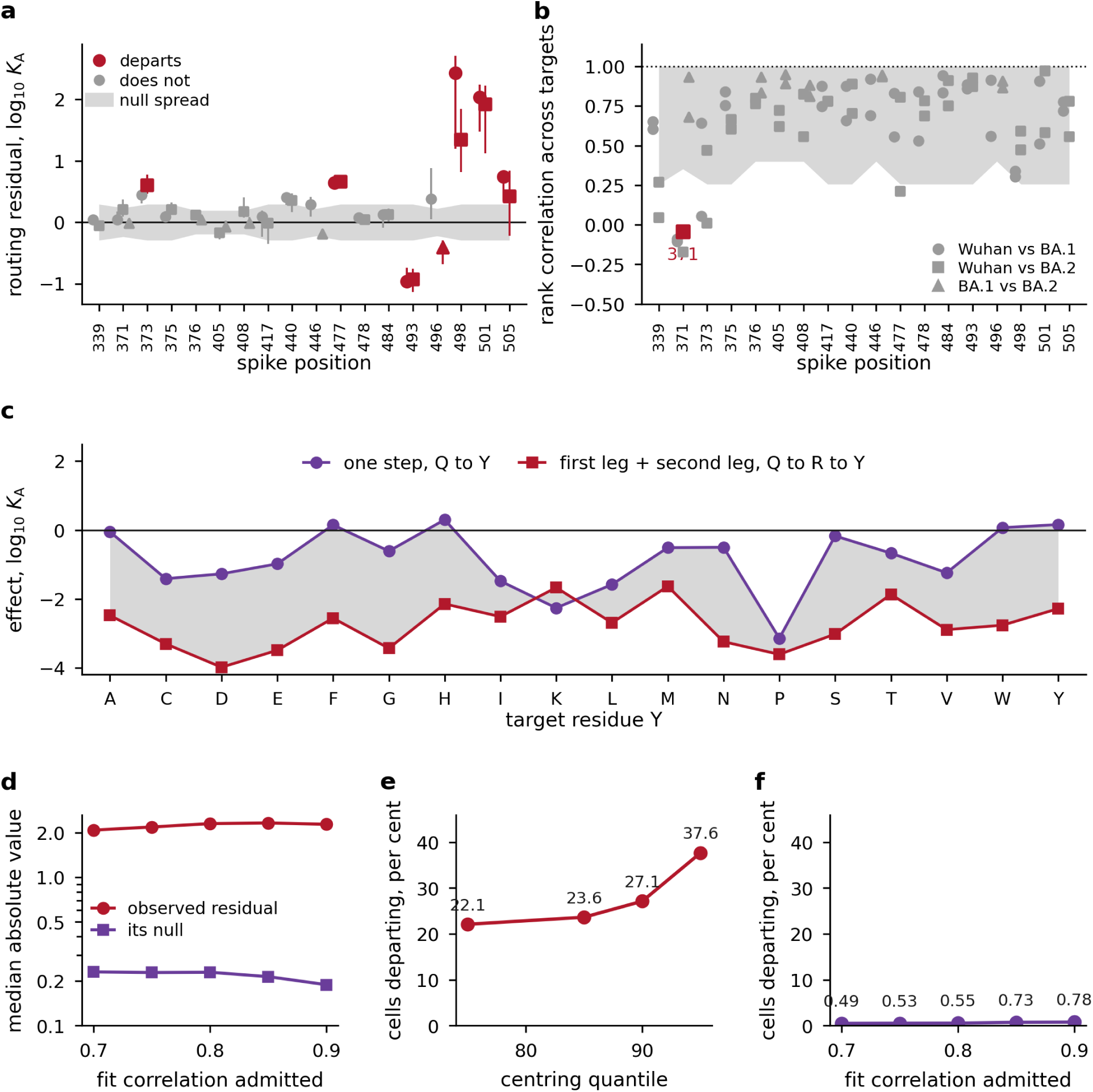
The one domain whose scale was checked, and the null that cannot be built anywhere else. (a) Routing residual by spike position for ACE2 binding, with the matched null for each position. Positions departing at a Benjamini-Hochberg adjusted p below 0.05 are marked. (b) Rank correlation between the one-step vector and the second-leg vector at the same positions, with the median of its matched null. The position that departs here is not among those that depart in (a). (c) Effects on binding for every target residue at the one position whose parental residue differs in all three backgrounds, measured from each parent, which is the reordering the rank statistic detects and the magnitude statistic does not. (d) Median absolute routing residual and the median of its null across the census, against the fit correlation at which a deposit pair is admitted after affine calibration. The two do not converge at any threshold. (e) Share of positions departing against the quantile used to estimate each deposit’s offset, for the estimated-centring correction. The answer moves with the estimate. (f) Share departing against the admitted fit correlation, for the magnitude statistic and for the rank statistic. The rank statistic is flat, as an increasing affine map cannot move it.

### Why the magnitude statistic has no null outside this domain

Outside this domain the same magnitude statistic has no null to be read against, and the reason is worth setting down because it is the second place a null fails here. Write a deposit’s reported effect as an affine function of the underlying one, s = a.e + c, where a is the scale the depositing laboratory used and c its offset. For a position whose first background carries X and whose second carries Z, the routing residual is the one-step effect minus the sum of the two legs, and under exact composition it reduces to

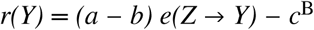

so the residual is not zero under composition unless the two deposits share both their scale and their offset. The offset of the second deposit survives alone and uncancelled. No cost in a phenotype unit is therefore reported outside this domain, and closing that needs a deposit set built for it, with a third background differing at a position already shown to compose, or one background scanned under two parentages.

Applying the receptor binding domain null across the census without correction returns 79.7 per cent of 2,954 cells departing, with a median first-leg effect of +4.8178 and a negative median residual in 93.7 per cent of them. A first leg improving a phenotype by nearly five log units at the median position is a property of the deposits and not of the proteins. Estimating the offset from a quantile of each deposit’s effect distribution removes those symptoms and replaces them with a dependence on the quantile, which moves the departure rate from 22.1 to 37.6 per cent across the swept range (Figure 5e).

### What calibration recovers, and what it does not

Fitting the map directly is the stronger correction, and for each pair of deposits we fit the least-squares affine map carrying one onto the other, using only the positions whose parental residue is the same in both, which are disjoint from the positions the test examines, and admit a pair when the fit’s correlation reaches a threshold that is swept. The anchors separate cleanly: the 6 same-phenotype receptor binding domain pairs fit between 0.860 and 0.965, and the 9 cross-phenotype pairs between 0.548 and 0.705 (Supplementary Table S3). At an admitted correlation of 0.80 the median absolute residual is 2.3022 against a null median of 0.2291 (Supplementary Table S2), and 1,863 of 1,942 cells exceed their own null (Figure 5d, 6f, Supplementary Table S5), and every cell of the run is listed in Supplementary Table S14.

That failure is a property of the deposits, because the null is the scatter of two deposits about the fitted line, which sets one measurement against one measurement, while the residual is a three-term combination of two measurements in the first deposit and one rescaled measurement in the second. No position whose parental residue is shared can supply the matched quantity, because forming the three-term combination requires a parent that differs between the deposits, and a differing parent is what defines a test position. The positions available to calibrate the statistic are the positions at which it cannot be formed. Fitting the map also relocates the surviving offset from the second deposit to the first instead of removing it.

### Where the departing positions sit

In the receptor binding domain the per-position rank test departs at 1 of 74 cells, at spike 371, and that position is not among the 9 the magnitude statistic flags. The two sets do not overlap (Figure 5b). The two statistics measure orthogonal components: a uniform penalty on the second step moves the magnitude of the residual and leaves the ordering of targets intact, while a change in which substitutions the position prefers moves the ordering and can leave the mean unchanged. The receptor binding domain signal is a shift, and the one position that reorders is spike 371, the only position whose parental residue differs in all three backgrounds (Figure 5c).

Where the departures sit is not arbitrary, and mapping the tested positions onto the deposited Omicron BA.1 receptor binding domain in complex with ACE2 [52], with the chain and the numbering offset solved for, not assumed, 198 of 201 residues of the deposited target match chain A at an offset of −3. The 7 positions departing on binding sit a median 3.5 Å from the receptor against 8.9 Å for the 11 that do not, and 6 of the 7 lie within 5 Å of it while none of the 11 do, at an exact p of 0.002 over every way of splitting the tested positions into a departing set of that size (Figure 6a, 7b).

**Figure 6.**
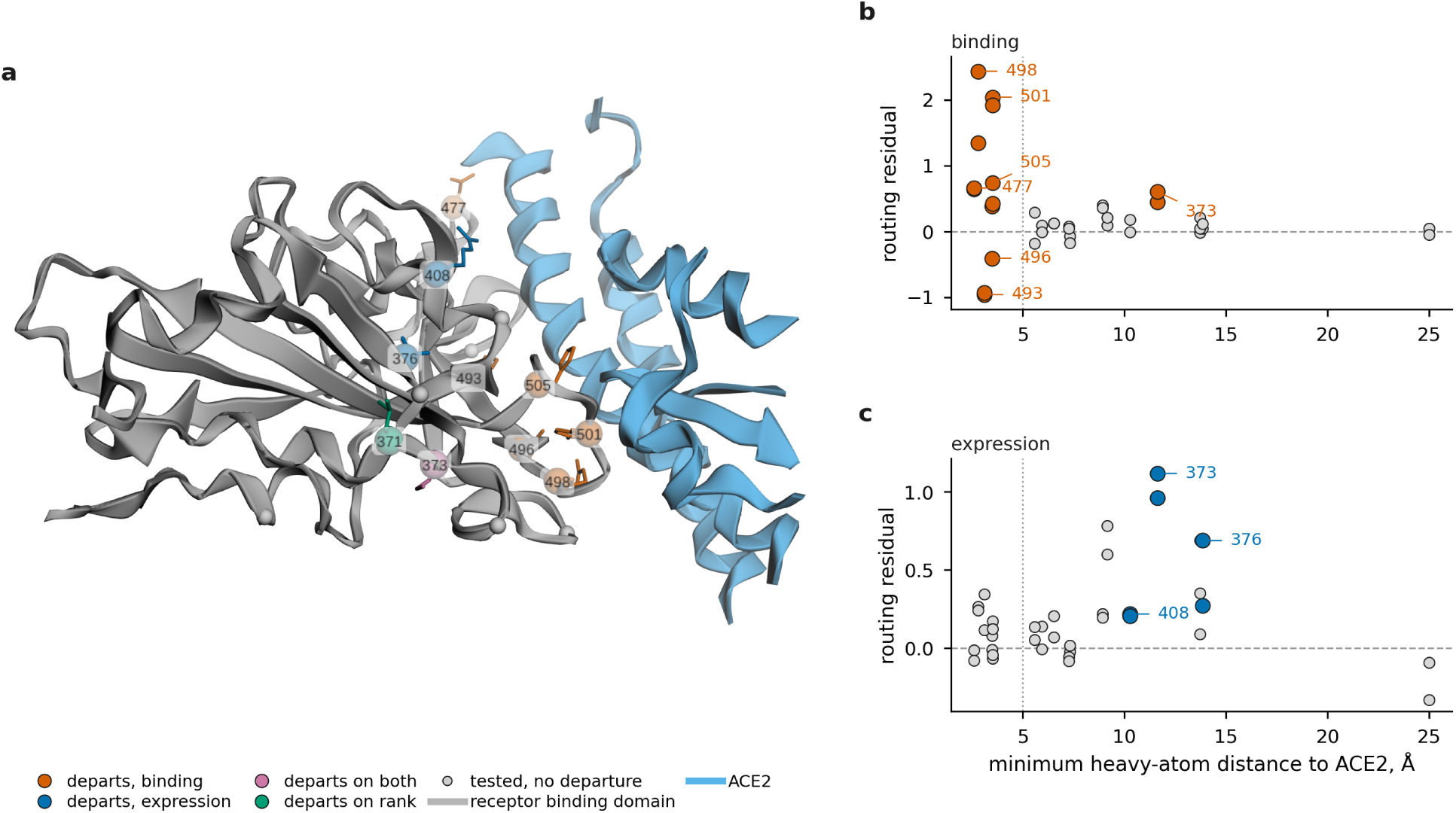
Where the departing positions sit, one phenotype at a time. (a) The tested positions on the deposited Omicron BA.1 receptor binding domain in complex with ACE2, with the chain and the numbering offset solved for, not assumed. Side chains are drawn for the departing positions and only the stretches of the receptor facing the domain are shown. (b) Routing residual against the minimum heavy-atom distance from the position to the receptor, for ACE2 binding, with the departing positions filled and labelled and the 5 Å contact distance marked. (c) The same for surface expression. The two phenotypes run in opposite directions, and the p values are exact over every split of the tested positions into a departing set of the observed size.

The expression phenotype runs the other way, with its 3 departing positions sitting a median 11.6 Å from the receptor against 5.9 Å for the rest, at an exact p of 0.020 (Figure 6c), and the position that reorders on the rank statistic sits 13.7 Å away. The two phenotypes are therefore not reporting one thing. Routing fails at the receptor contact surface for binding and away from it for expression, which is what a mechanism of interface entrenchment predicts and what a scale artefact common to the deposits does not. Pooling the phenotypes returns a weak interface effect and conceals the reversal, which is why they are kept apart throughout. The localisation was not specified before the departing positions were known, so it is reported as an observation over the 18 tested positions of this domain that a mechanism of entrenchment predicts.

### Routes through alanine

Alanine is the intermediate a design method reaches for most often, and 12,230 of the deduplicated routes pass through it. An intermediate earns that place by sitting one base change from each end of an exchange the code separates by two. Profiling the mutational landscape of the deposited alpha-helical transmembrane entries placed the valine to threonine exchange through an alanine intermediate and the glycine to asparagine exchange through a serine intermediate, lowering a barrier that two sequential base changes would otherwise impose [2], and a kernel-based analysis of toll-like receptor 5 in Epinephelus coioides indicated that the threonine to valine changes separating its membrane-bound and soluble forms may have been supported by alanine intermediates [3]. Serine and threonine may serve the same role between basic and acidic residues [4]. Each of those intermediates was inferred from sequence, and the deposited record measures what such a route costs where the scans carry it.

Across systems the shift correlates with the fraction of a system’s positions routed through alanine at 0.236, so systems in which more routes pass through alanine depart less (Supplementary Table S12). The sub-claim under test is whether a route through alanine composes as well as a route through another residue, on these deposits and on this rank statistic, and the correlation runs across systems and not within one.

## Conclusions

Deep mutational scanning has made the first step of a substitution path routine, and the second step is already deposited wherever two scans of one protein differ by a single parental substitution. What has been missing is the arithmetic that says when two such deposits can be added. Expanding a route into its two legs supplies it, because the discrepancy between a route and its legs is epistasis between the tested position and wherever else the two parents differ, and it vanishes when they differ nowhere else. The comparison of two deposits therefore has a value fixed in advance, and the census assembles 102,421 such routes over 65 protein clusters from deposits that already exist. Over the 741 pairs the record admits the median is 0.965, so the arithmetic holds wherever the test applies.

Benchmarking a variant effect predictor against a single deposit cannot exceed the agreement two deposits of one protein reach at the positions both report, a ceiling of 0.80 at a position drawn at random, so a benchmark that scores 55 predictors against 26 proteins [15] works against a bound belonging to the data and not to the predictors. Functional evidence entering clinical variant interpretation under the PS3 and BS3 criteria [17] carries the same bound, and imputing the scores a deposit does not report [19] presumes a commensurability the test settles pair by pair before the imputation runs. The null these comparisons are ordinarily read against returns +0.227 where zero is required, so a routing shift smaller than that bias is not evidence of a cost. None of this is recoverable from the deposited metadata, where the pairs grouped under one experiment set are the ones that agree least, so each pair has to be cleared on its own.

Antibiotic resistance is reached through several substitutions whose order governs which alleles are available at all, and the census holds the TEM *β*-lactamases on backgrounds that single substitutions define, deposited by successive studies along one adaptive path [40,41,42,43,44]. Those are the scans a reader would reach for first, and they return a median of 0.25 where one is required, so a resistance path read from two of them is the case the test has to clear before reachability is claimed. Antigenic escape is the same problem in another setting: the receptor binding domain is deposited on the Wuhan Hu-1, BA.1 and BA.2 backgrounds [49,50], an entrenchment claim at Q498R and N501Y rests on reading those deposits against one another, and the positions whose routes depart cluster on the receptor contact surface for binding and off it for expression, so each such comparison can be checked position by position before a path through an intermediate is believed. The paths that matter are assembled across experiments, and that is where nothing in the record vouches for the addition.

Methods that design a path through an intermediate gain most directly, since they can take the second leg from a deposit whose addition to the first has been certified instead of assumed, and alanine, the intermediate such methods reach for most often, is the one 12,230 of the assembled routes pass through. The pairs that admit the test today come mostly from one proteolysis assay applied at scale [38], and every further library of that kind adds pairs the test can certify at one rank correlation per position. The combinatorially complete library of the fifteen substitutions separating Wuhan Hu-1 from BA.1 [27] already carries paths inside one experiment, and extending that design to two substitutions at a single position would deliver a route end to end; exactly 1 of the 796 qualifying pairs in the record is of that form today, which makes it an inexpensive experiment whose answer is known in advance. The deposited record has been read as a catalogue of single substitutions, and the arithmetic given here turns it into a map of the paths between them. Resistance, antigenic escape and protein design all depend on paths of that kind, and the test certifies each one before it is read.

## Methods

### Deposits and the route census

Score sets and target sequences were retrieved from the MaveDB API [32] on 11 November 2026. Coding sequences were translated and protein sequences taken as deposited. A target was set aside when it carried no sequence, when a coding length was not a multiple of three, when the translation carried an internal stop, or when it was shorter than twenty residues. Two targets of equal length differing at one or more positions by at most 25.0 per cent of their residues form a background pair, and positions correspond by index with no alignment step.

At a position where the first background carries X and the second carries Z, a route to target residue Y requires the one-step effect of X to Y and the first leg of X to Z in the first scan, and the second leg of Z to Y in the second. Y runs over the residues other than X and Z. Synonymous parental entries and stop codons are excluded. Effects are read from the per-variant mean where a deposit reports one per library and from the deposited score otherwise, a distinction that matters because reading the per-library column of a two-library deposit takes each variant twice at a single library’s value.

Reference residues in the deposited variant names are checked against the translated target at every position used, and a disagreement is a fatal condition and not a warning. Redundant deposits of one target are collapsed onto distinct sequences before any count is reported.

### The magnitude statistic and its censoring

The routing residual is the one-step effect minus the sum of the two legs, and the cost of the intermediate is the first leg, reported beside the residual and never folded into it. For the receptor binding domain the null is the discrepancy between the two scans at positions whose parental residue is identical in both, where the same substitution is measured twice and no routing is involved.

The statistic is the median absolute residual at a position, referred to the median absolute discrepancy over 10,000 random draws of the same number of control pairs from the same background pair and phenotype, drawn without replacement, with p the one-sided upper-tail proportion. Multiplicity is controlled by the Benjamini-Hochberg procedure [53] within phenotype, and results are never pooled across phenotypes.

Only the Wuhan binding scan censors, at −4.8 with 101 unique substitutions at that value, and the floor detected in each deposit is reported in Supplementary Table S6. A route is excluded when any of its three measurements lies at or below the scan’s censoring threshold plus a margin, and the margin is swept over 0.00, 0.01, 0.05, 0.10, 0.20, 0.30, 0.50 with every table reported at each value (Supplementary Table S11).

### Base changes between residues

Codon distances use the standard genetic code. The distance between two amino acids is the smallest number of positions at which a codon of the first and a codon of the second differ, minimised over both sets of synonymous codons, and stop codons are excluded. For each route fan, defined by a deposit pair and a position at which a complete route exists, every target residue other than the parental residue X and the intermediate Z contributes one target slot, and a slot is counted once per distinct combination of the two deposit identifiers and the position so that redundant deposits of one target do not inflate the tally.

### The rank statistic

Under composition the one-step vector and the second-leg vector over the target residues at one position are related by an affine map, because the first leg does not depend on Y, so their ranks agree. The statistic is the Spearman rank correlation [54] between them with ties averaged, which is unchanged by any increasing affine rescaling of either deposit and therefore requires no estimate of a deposit’s offset or scale.

The null is the same statistic at the positions whose parental residue is identical in both deposits, where the two vectors are one quantity measured twice. A draw selects such a position uniformly from the same deposit pair and, where it carries more target residues than the test position, a random subset of them matched to the test position’s count. The reference is 10,000 draws and p is the one-sided lower-tail proportion. Size was checked by drawing values from the null and testing them against a reference built from disjoint draws.

The census lists every deposit pair in both directions, since a route has a direction, and the rank correlation is symmetric in the two deposits. Counts of cells and of comparisons are therefore collapsed onto unordered pairs before they are reported, and counts of routes are not.

### The exact test, and the pairs that admit it

A deposit pair admits the test when both target sequences translate, have equal length, differ at exactly one position and that position is the one the census row tests. The condition is checked on the translated sequences and not on the deposit names. At such a pair the one-step vector and the second-leg vector are built over the target residues measured in both deposits, excluding the two parental residues, and a pair is admitted when at least 6 residues remain. The statistic is the same Spearman correlation as above and the predicted value is one, so no null is drawn and no multiplicity arises.

The control at the same pair is that statistic at the positions whose parental residue the two deposits share, each matched to the test position on the number of target residues by a seeded random subset, with the median taken over at most 40 such positions. Amino acids are iterated in sorted order so that a seeded subset is the same on every run, and both scripts were run twice and their outputs compared byte for byte.

### The nulls and the trend against background separation

The dispersion-matched null differs from that control in one respect. Each candidate control position carries the standard deviation of the effects it contributes in each deposit, and the 10 positions nearest the test position in the plane of those two logarithms are taken instead of 10 drawn at random. The two nulls are computed from one candidate pool in one pass, so they differ only in which positions they select.

The trend against background separation is computed over every deposit pair in the census, not only over those admitting the test. The number of separating positions is counted on the translated sequences, at most 12 test positions are taken per pair when more exist, and the pair’s shift is the mean over its test positions of the test statistic minus the mean over the same positions of the null. Bins are reported as medians, and the correlation against the logarithm of the separation is a product-moment one with the test position count partialled out.

### Assay identity and the reliability bound

Two deposits are counted as one assay when they sit in one MaveDB experiment, which is a curated fact, or when their score set titles agree once the target’s own name and any parenthesised content are removed, which catches one study depositing many targets under one assay. Every other pair is counted as a different assay, and the rule is mechanical so that a pair is not reclassified after its statistic is seen.

The reliability bound follows the classical attenuation relation [54]. Two deposits of one protein at a position whose parental residue they share are treated as parallel measurements of one quantity with independent error, so the correlation between either and the quantity is at most the square root of the correlation between them. The relation is exact for the product-moment correlation and is applied here to the rank correlation as an approximation, and it assumes what the test of the same pairs establishes, that the two deposits measure one quantity at all.

### Scale calibration and system kind

For each pair of deposits the least-squares affine map carrying the second onto the first was fitted using only the positions whose parental residue is shared, which are disjoint from the positions the test examines, so a fitted map cannot carry information from a test position into its own calibration. A pair was admitted when the fit’s Pearson correlation reached a threshold, swept over 0.70, 0.75, 0.80, 0.85, 0.90.

A protein system is the set of deposit pairs sharing a base target name. A deposit name carrying a parenthesised point substitution or design tag marks a designed background. The rule is applied to the deposited names and not to a judgement about each system, and both groups are reported.

The census holds every deposit pair in both directions, and once for each redundant pairing of the deposits that realise the same two target sequences, so the cells are collapsed to one per distinct pair of target sequences and position, keeping the cell with the most shared target residues. Collapsing takes the 47,607 cells the census offers to 4,439, and a system is tested when at least 8 survive.

### The structure and the distances to the receptor

Positions of the receptor binding domain are placed on the deposited Omicron BA.1 domain in complex with ACE2 [52], Protein Data Bank entry 7WPB. Neither the chain nor the numbering offset is assumed: the deposited target sequence is translated and matched against every chain at every offset from minus thirty to plus thirty residues, and the best match is taken, which is chain A at an offset of −3 with 198 of 201 residues agreeing. A match below nine residues in ten is a fatal error, not a warning. The remaining chain is the receptor, and the distance reported for a position is the smallest distance from any of its heavy atoms to any heavy atom of that chain, with hydrogens and alternate locations beyond the first discarded.

Whether the departing positions sit closer to the receptor than the positions that do not is tested exactly and separately within each phenotype. The Mann-Whitney rank sum [55] of the departing set is compared against its value under every way of splitting the tested positions into a departing set of the observed size, with ties averaged, and the one-sided p is the proportion of splits whose rank sum is at most the observed one. With 18 positions this enumerates 31,824 splits, so no distributional approximation enters and no external statistics package is used.

## Data availability

All measurements analysed here are previously deposited and were retrieved from MaveDB [32]. The score sets underlying the receptor binding domain analysis are urn:mavedb:00000044-a-1 and urn:mavedb: 00000044-b-2 for the Wuhan Hu-1 background [49], urn:mavedb:00000668-a-1 and urn:mavedb:00000669a-1 for BA.1, and urn:mavedb:00000670-a-1 and urn:mavedb:00000671-a-1 for BA.2 [50]. The census are listed with their checksums in Supplementary Table S8. The census reads 688 score sets in total, and the identifier of every one of them is carried in the route census table of the accompanying repository, one row per deposit pair and position. The supplementary information holds the per-position detail behind the results reported here. The structure is Protein Data Bank entry 7WPB [52].

## Code availability

The analysis scripts, the derived tables and the figure sources are at https://github.com/karagol-taner/twostep-substitution-routes.

## Author contributions

T.K. and A.K. contributed equally to this work. T.K. conceived the study, designed and implemented the analysis and wrote the manuscript. A.K. contributed to the design of the statistical controls and to the manuscript. Both authors read and approved the final version.

## Competing interests

T.K. and A.K. are named co-inventors (50 percent / 50 percent share) on Turkish patent applications concerning STING and VAMP2 and a pH-dependent protein engineering methodology. A.K. is a Novo Nordisk AI Fellow at MIT. T.K. holds an award from the Anthropic AI for Science program for a separate project at MIT. Authors declare no competing interests related to the material presented in this study.

## Funding

No grant funding was received for this work. The large language model assistance declared below used model credits from an Anthropic AI for Science award held by T.K. for unrelated work, and Anthropic had no role in the design of the study, the analysis, the interpretation of the results or the decision to publish. No other interest declared above provided financial or operational support.

## Ethics approval

Ethics approval was not required for this computational study, which did not involve animal subjects, human participants or identifiable data.

## Declaration of Generative AI and AI-Assisted Technologies in the Manuscript Preparation Process

During the preparation of this work, the authors used Claude Opus 5 (Anthropic) for drafting and for streamlining analysis code. The authors reviewed and edited the output as needed and take full responsibility for the content of the published article.

## Notes

https://github.com/karagol-taner/twostep--substitution-routes

