## Supplementary Information for "Most substitutions need more than one base change and an exact test tells which deposited scans measure the route"

### Table of contents


| | Item | Where |
|---|---|---|
| 1. | Supplementary Table S1. Positions of the receptor binding domain departing from composition in the magnitude of the routing residual, at a censoring margin of zero. | Pg. 3 |
| 2. | Supplementary Table S2. Every table of the calibrated run at every admitted fit correlation, as the calibration specification fixed before the run. | Pg. 4 |
| 3. | Supplementary Table S3. The 15 affine fits among the six receptor binding domain deposits, which anchor the admission threshold. | Pg. 5 |
| 4. | Supplementary Table S4. The commensurability check on the receptor binding domain deposits, which is the one place in the census where a scale was checked rather than assumed. | Pg. 6 |
| 5. | Supplementary Table S5. The calibrated run one protein system at a time, at an admitted fit correlation of 0.80. | Pg. 7 |
| 6. | Supplementary Table S6. Censoring in the six receptor binding domain deposits, detected from the deposited scores rather than taken from a published limit of detection. | Pg. 8 |
| 7. | Supplementary Table S7. The shift between the test position and its null by the number of positions separating the two backgrounds, under both nulls. | Pg. 9 |
| 8. | Supplementary Table S8. The six score sets the receptor binding domain analysis reads, as retrieved. | Pg. 10 |
| 9. | Supplementary Table S9. The three rank and product-moment statistics over the same 741 deposit pairs, at the test position and at the control positions. | Pg. 11 |
| 10. | Supplementary Table S10. How the deposited record relates the two deposits of a pair, for every pair and for the pairs the test admits. | Pg. 12 |
| 11. | Supplementary Table S11. The censoring margin sweep over the six receptor binding domain deposits, fixed in the analysis plan before the residual was computed. | Pg. 13 |
| 12. | Supplementary Table S12. The combined within-system rank test, calibrated, one protein system at a time. | Pg. 14 |
| 13. | Supplementary Table S13. Every deposit pair the exact test admits, one row per pair. | Supplementary_Tables.xlsx |
| 14. | Supplementary Table S14. Every cell of the calibrated run at an admitted fit correlation of 0.80, one row per cell. | Supplementary_Tables.xlsx |

### Supplementary tables

**Supplementary Table S1. Positions of the receptor binding domain departing from composition in the magnitude of the routing residual, at a censoring margin of zero.**

| Spike[1] | Phenotype | Pair | X → Z[2] | Routes | Median residual[3] | Null median | p (BH)[4] |
|---|---:|---:|---:|---:|---:|---:|---:|
| 493 | binding | Wuhan / BA.1 | Q → R | 18 | −0.9600 | 0.323 | 0.00062 |
| 498 | binding | Wuhan / BA.1 | Q → R | 18 | +2.4342 | 0.323 | 0.00062 |
| 501 | binding | Wuhan / BA.1 | N → Y | 18 | +2.0405 | 0.323 | 0.00062 |
| 493 | binding | Wuhan / BA.2 | Q → R | 18 | −0.9315 | 0.263 | 0.00062 |
| 498 | binding | Wuhan / BA.2 | Q → R | 18 | +1.3429 | 0.263 | 0.00062 |
| 501 | binding | Wuhan / BA.2 | N → Y | 18 | +1.9217 | 0.263 | 0.00062 |
| 505 | binding | Wuhan / BA.2 | Y → H | 18 | +0.4207 | 0.263 | 0.00106 |
| 496 | binding | BA.1 / BA.2 | S → G | 18 | −0.4082 | 0.123 | 0.00185 |
| 376 | expression | BA.1 / BA.2 | T → A | 18 | +0.2706 | 0.121 | 0.00185 |
| 373 | expression | Wuhan / BA.2 | S → P | 18 | +1.1177 | 0.309 | 0.00185 |
| 477 | binding | Wuhan / BA.2 | S → N | 18 | +0.6568 | 0.263 | 0.00411 |
| 505 | binding | Wuhan / BA.1 | Y → H | 18 | +0.7399 | 0.323 | 0.00481 |
| 373 | binding | Wuhan / BA.2 | S → P | 18 | +0.6007 | 0.263 | 0.00706 |
| 408 | expression | BA.1 / BA.2 | R → S | 18 | +0.2172 | 0.121 | 0.02590 |
| 477 | binding | Wuhan / BA.1 | S → N | 18 | +0.6437 | 0.323 | 0.02713 |

[1]Spike numbering, obtained by solving for the offset between the deposited numbering and spike numbering rather than assuming it.

[2]X is the parental residue of the first background at that position and Z the parental residue of the second, so the first leg is X to Z and the second leg runs from Z.

[3]The one-step effect minus the sum of the two legs, in $\log_{10} K_a$ for binding and $\log_{10}$ expression for expression. A positive value means the two-step sum predicts a worse outcome than the direct substitution delivers.

[4]Benjamini-Hochberg adjusted within phenotype over the 37 positions tested on each.

**Supplementary Table S2. Every table of the calibrated run at every admitted fit correlation, as the calibration specification fixed before the run.**

| Threshold[1] | Deposit pairs | Cells[2] | Systems | Departing, magnitude[3] | Departing, rank[3] | Median \|residual\|[4] | Null median[4] | Median ρ | Null median ρ |
|---|---|---|---|---|---|---|---|---|---|
| 0.70 | 7,669 | 4,859 | 56 | 4,265 | 24 | 2.082 | 0.231 | 0.732 | 0.645 |
| 0.75 | 7,059 | 4,349 | 49 | 3,939 | 23 | 2.184 | 0.228 | 0.748 | 0.680 |
| 0.80 | 6,232 | 3,842 | 39 | 3,588 | 21 | 2.302 | 0.229 | 0.767 | 0.733 |
| 0.85 | 4,942 | 3,164 | 39 | 2,997 | 23 | 2.326 | 0.214 | 0.769 | 0.783 |
| 0.90 | 3,178 | 2,191 | 31 | 2,078 | 17 | 2.278 | 0.189 | 0.771 | 0.819 |

[1]The Pearson correlation the affine map fitted on shared-parent positions must reach before a deposit pair is admitted. The specification swept 0.70 to 0.90 and required every table at every value.

[2]Collapsed onto one cell per distinct pair of target sequences and position.

[3]Cells whose statistic departs at a Benjamini-Hochberg adjusted p below 0.05.

[4]The magnitude statistic is the median absolute routing residual at a position, in the units each deposit reports.

**Supplementary Table S3. The 15 affine fits among the six receptor binding domain deposits, which anchor the admission threshold.**

| From deposit | Phenotype | Onto deposit | Phenotype | Same phenotype | Matched substitutions[1] | Fit correlation[2] |
|---|---|---|---|---|---|---|
| 00000668-a-1 | binding | 00000670-a-1 | binding | yes | 3,687 | 0.965 |
| 00000669-a-1 | expression | 00000671-a-1 | expression | yes | 3,687 | 0.954 |
| 00000044-b-2 | expression | 00000671-a-1 | expression | yes | 3,476 | 0.906 |
| 00000044-b-2 | expression | 00000669-a-1 | expression | yes | 3,513 | 0.878 |
| 00000044-a-1 | binding | 00000668-a-1 | binding | yes | 3,517 | 0.863 |
| 00000044-a-1 | binding | 00000670-a-1 | binding | yes | 3,480 | 0.860 |
| 00000044-b-2 | expression | 00000670-a-1 | binding | no | 3,476 | 0.705 |
| 00000044-b-2 | expression | 00000668-a-1 | binding | no | 3,513 | 0.694 |
| 00000670-a-1 | binding | 00000671-a-1 | expression | no | 3,801 | 0.651 |
| 00000044-a-1 | binding | 00000044-b-2 | expression | no | 3,797 | 0.643 |
| 00000668-a-1 | binding | 00000671-a-1 | expression | no | 3,687 | 0.623 |
| 00000669-a-1 | expression | 00000670-a-1 | binding | no | 3,687 | 0.621 |
| 00000668-a-1 | binding | 00000669-a-1 | expression | no | 3,819 | 0.620 |
| 00000044-a-1 | binding | 00000671-a-1 | expression | no | 3,480 | 0.569 |
| 00000044-a-1 | binding | 00000669-a-1 | expression | no | 3,517 | 0.548 |

[1]Substitutions measured in both deposits at positions whose parental residue they share, which are the positions the fit uses and are disjoint from the positions the test examines. A pair is fitted only when at least 100 are available.

[2]The six same-phenotype fits run from 0.860 to 0.965 and the nine cross-phenotype fits from 0.548 to 0.705, so any threshold from 0.70 to 0.85 admits the first group and rejects the second.

**Supplementary Table S4. The commensurability check on the receptor binding domain deposits, which is the one place in the census where a scale was checked rather than assumed.**

| Background | Phenotype | Column | Rows compared | Parental absolute | Largest deviation[1] | Tolerance | Relation |
|---|---|---|---|---|---|---|---|
| BA.1 | binding | bind | 3,819 | 9.222 | 0.017 | 0.005 | offset varies |
| BA.1 | expression | expr | 3,819 | 9.732 | 0.000 | 0.005 | constant offset |
| BA.2 | binding | bind | 4,002 | 9.255 | 0.017 | 0.005 | offset varies |
| BA.2 | expression | expr | 4,002 | 9.854 | 0.026 | 0.005 | offset varies |

[1]The largest departure of any row from the median offset between the deposited absolute score and the parental score plus the reported difference. A constant offset means the deposited absolute is the parental value plus the difference, to within the deposited rounding.

**Supplementary Table S5. The calibrated run one protein system at a time, at an admitted fit correlation of 0.80.**

| System[1] | Cells[2] | Departing, magnitude | Departing, rank | Median \|residual\| | Null median | Median ρ | Null median ρ |
|---|---|---|---|---|---|---|---|
| protein G | 1,337 | 1,288 | 12 | 2.445 | 0.258 | 0.868 | 0.847 |
| ice-structuring protein | 1,006 | 1,003 | 0 | 2.325 | 0.197 | 0.633 | 0.585 |
| SPTAN1 | 387 | 358 | 0 | 1.749 | 0.268 | 0.825 | 0.845 |
| spa | 180 | 164 | 1 | 2.944 | 0.229 | 0.716 | 0.640 |
| HPLC 12 | 139 | 139 | 0 | 2.354 | 0.214 | 0.615 | 0.612 |
| affibody | 132 | 122 | 1 | 3.030 | 0.161 | 0.685 | 0.605 |
| ubiquitin | 130 | 125 | 2 | 1.989 | 0.239 | 0.707 | 0.769 |
| rplI | 95 | 70 | 0 | 2.737 | 0.241 | 0.953 | 0.892 |
| SARS-CoV-2 RBD | 74 | 21 | 0 | 0.222 | 0.261 | 0.841 | 0.814 |
| HPLC 12 isoform | 55 | 55 | 0 | 2.374 | 0.183 | 0.541 | 0.559 |
| U14Ub1 | 37 | 37 | 0 | 2.766 | 0.277 | 0.787 | 0.854 |
| zinc finger | 32 | 4 | 0 | 0.093 | 0.114 | 0.436 | 0.361 |
| FYN SH3 domain | 28 | 27 | 0 | 1.802 | 0.137 | 0.926 | 0.918 |
| Protein LG | 28 | 24 | 2 | 1.711 | 0.251 | 0.902 | 0.854 |
| Spectrin alpha, non-erythroc | 27 | 27 | 0 | 5.125 | 0.229 | 0.820 | 0.779 |
| homeobox domain | 16 | 2 | 0 | 0.092 | 0.104 | 0.506 | 0.546 |
| StoA | 13 | 10 | 0 | 1.834 | 0.237 | 0.891 | 0.831 |
| StoC | 13 | 12 | 0 | 1.575 | 0.181 | 0.891 | 0.833 |
| entD | 12 | 11 | 0 | 0.964 | 0.131 | 0.866 | 0.908 |
| influenza NP | 8 | 8 | 0 | 3.516 | 0.262 | 0.995 | 0.876 |
| NfeD domain-containing prote | 8 | 7 | 0 | 2.238 | 0.159 | 0.958 | 0.620 |
| Uncharacterized protein MJ11 | 8 | 7 | 0 | 1.475 | 0.181 | 0.986 | 0.822 |
| cspLA | 8 | 8 | 0 | 1.893 | 0.210 | 0.832 | 0.787 |
| cylR2 | 8 | 8 | 0 | 1.659 | 0.158 | 0.902 | 0.856 |
| SRC | 8 | 8 | 0 | 3.463 | 0.203 | 0.968 | 0.868 |

[1]A system is the set of deposit pairs sharing a base target name, which groups engineered variants of one protein with its wild type and is not a clade, a family or a fold.

[2]Collapsed onto one cell per distinct pair of target sequences and position. The 39 systems here are those surviving the admission threshold with at least the required number of cells.

**Supplementary Table S6. Censoring in the six receptor binding domain deposits, detected from the deposited scores rather than taken from a published limit of detection.**

| Deposit | Background | Phenotype | Substitutions measured | Lowest score | Detected floor[1] | Substitutions at the floor |
|---|---|---|---|---|---|---|
| 00000668-a-1 | BA.1 | binding | 3,819 | −3.922 | none | 0 |
| 00000669-a-1 | BA.1 | expression | 3,819 | −1.996 | none | 0 |
| 00000670-a-1 | BA.2 | binding | 4,002 | −3.925 | none | 0 |
| 00000671-a-1 | BA.2 | expression | 4,002 | −2.139 | none | 0 |
| 00000044-a-1 | Wuhan | binding | 3,802 | −4.840 | −4.800 | 101 |
| 00000044-b-2 | Wuhan | expression | 3,798 | −4.070 | none | 0 |

[1]A score is called a floor when the same value repeats at least 20 times in the lower tail. The exclusion margin above any detected floor was swept over 0.00, 0.01, 0.05, 0.10, 0.20, 0.30, 0.50 and every result in the manuscript is reported at a margin of zero.

**Supplementary Table S7. The shift between the test position and its null by the number of positions separating the two backgrounds, under both nulls.**

| Positions differing | Deposit pairs | Median shift, matched on count[1] | Median shift, matched on dispersion[1] |
|---|---:|---:|---:|
| 1 | 741 | +0.215 | +0.107 |
| 2 | 1,043 | −0.055 | −0.046 |
| 3 | 576 | −0.091 | −0.023 |
| 4-6 | 849 | −0.092 | −0.056 |
| 7-20 | 1,553 | −0.092 | −0.057 |
| >20 | 5 | −0.126 | −0.116 |

[1]The algebra fixes this at zero in the first row and leaves it unknown in every other. The first row is therefore the only one whose value can be checked, and it is the row that carries the whole correlation against separation.

**Supplementary Table S8. The six score sets the receptor binding domain analysis reads, as retrieved.**

| **Score set** | **Background** | **Phenotype** | **Variants declared** | **Rows read[1]** | **MD5 of the file read** |
|---|---|---|---|---|---|
| 00000044-a-1 | Wuhan | binding | 8,040 | 8,040 | 34b61ccd4e408779ad0e447052e15bdd |
| 00000044-b-2 | Wuhan | expression | 8,040 | 8,040 | e0101efe13d31aeb62010803bad158ed |
| 00000668-a-1 | BA.1 | binding | 3,819 | 3,819 | 23e77d97df94b0234dedfa65e9e54f8f |
| 00000669-a-1 | BA.1 | expression | 3,819 | 3,819 | e463aaeab2d6534a15e260081302fc3b |
| 00000670-a-1 | BA.2 | binding | 4,020 | 4,020 | d49e4b6b6865698196ea6fef5d53699e |
| 00000671-a-1 | BA.2 | expression | 4,020 | 4,020 | e9dc5bef4ab3db85acb98979610b00e8 |

[1]Rows in the file retrieved from the MaveDB API. All six deposit the same target, the SARS-CoV-2 receptor binding domain, on the three parental backgrounds shown. The checksum is of the file the analysis read, so a rerun can establish that it read the same bytes.

**Supplementary Table S9. The three rank and product-moment statistics over the same 741 deposit pairs, at the test position and at the control positions.**

| Statistic | Median, test | Q1 | Q3 | Above 0.9, test | Median, control | Above 0.9, control |
|---|---|---|---|---|---|---|
| Spearman | 0.965 | 0.897 | 0.984 | 552 | 0.817 | 172 |
| Pearson | 0.984 | 0.944 | 0.993 | 607 | 0.817 | 174 |
| Kendall | 0.882 | 0.752 | 0.922 | 286 | 0.660 | 2 |

[1]Every one of the three is fixed at one by the same expansion, at the test position and at the control positions alike, so each column is read against one and not against the others.

**Supplementary Table S10. How the deposited record relates the two deposits of a pair, for every pair and for the pairs the test admits.**

| Relation in the deposited record | All deposit pairs | Pairs at one differing position |
|---|---:|---:|
| same experiment | 10 | 2 |
| same experiment set, different experiment | 104 | 92 |
| shared publication, different experiment set | 128 | 20 |
| unrelated deposits | 9,348 | 1,451 |

[1]Counted on undirected pairs. One experiment is the strongest relation the record carries, since it is curated, and it holds for 2 of the admitted pairs.

**Supplementary Table S11. The censoring margin sweep over the six receptor binding domain deposits, fixed in the analysis plan before the residual was computed.**

| Margin [1] | Binding routes | Binding usable | Binding excluded | Expression routes | Expression usable | Expression excluded |
|---|---|---|---|---|---|---|
| 0.00 | 666 | 666 | 0 | 666 | 666 | 0 |
| 0.01 | 666 | 665 | 1 | 666 | 666 | 0 |
| 0.05 | 666 | 664 | 2 | 666 | 666 | 0 |
| 0.10 | 666 | 664 | 2 | 666 | 666 | 0 |
| 0.20 | 666 | 660 | 6 | 666 | 666 | 0 |
| 0.30 | 666 | 658 | 8 | 666 | 666 | 0 |
| 0.50 | 666 | 655 | 11 | 666 | 666 | 0 |

[1]A route is excluded when any of its three measurements sits within this margin of the censoring floor of its own scan, in the units that scan reports. At a margin of zero no route is excluded, since only one of the six deposits censors at all.

**Supplementary Table S12. The combined within-system rank test, calibrated, one protein system at a time.**

| System[1] | Kind | Cells[2] | Cells via alanine | Deposit pairs | Mean ρ | Null mean ρ | Shift[3] | p[4] |
|---|---|---|---|---|---|---|---|---|
| UBB | designed background | 40 | 1 | 9 | 0.268 | 0.457 | −0.188 | 5.0e−3 |
| CYP2C19 | natural variant | 30 | 1 | 2 | 0.406 | 0.570 | −0.164 | 2.2e−2 |
| CYP2C9 | natural variant | 33 | 1 | 2 | 0.422 | 0.570 | −0.148 | 2.3e−2 |
| HPLC 12 isoform | designed background | 69 | 9 | 18 | 0.379 | 0.511 | −0.133 | 1.5e−3 |
| NP, H3N2 Aichi/1968 | natural variant | 30 | 1 | 1 | 0.242 | 0.366 | −0.124 | 1.6e−2 |
| NP | natural variant | 30 | 1 | 1 | 0.165 | 0.287 | −0.122 | 2.2e−2 |
| NP, H1N1 PR/1934 | natural variant | 30 | 3 | 1 | 0.165 | 0.286 | −0.121 | 2.2e−2 |
| Ice-structuring protein | designed background | 311 | 53 | 37 | 0.415 | 0.526 | −0.111 | 8.8e−4 |
| Type-3 ice-structuring protein HPLC 12 isoform | designed background | 70 | 9 | 18 | 0.417 | 0.527 | −0.110 | 5.0e−3 |
| HPLC 12 | designed background | 170 | 32 | 35 | 0.434 | 0.528 | −0.094 | 8.8e−4 |
| StoC | natural variant | 13 | 1 | 1 | 0.754 | 0.847 | −0.092 | 4.8e−2 |
| StoA | natural variant | 13 | 0 | 1 | 0.754 | 0.847 | −0.092 | 4.8e−2 |
| Ice-structuring protein RD1 | designed background | 368 | 42 | 33 | 0.347 | 0.434 | −0.087 | 8.8e−4 |
| Type-3 ice-structuring protein HPLC 12 | designed background | 717 | 124 | 144 | 0.438 | 0.519 | −0.082 | 8.8e−4 |
| SARS-CoV-2 Omicron BA.2 receptor binding domain | natural variant | 22 | 0 | 4 | 0.433 | 0.503 | −0.071 | 1.9e−1 |
| CHD4 Zinc finger, PHD-finger domain | natural variant | 12 | 1 | 3 | 0.228 | 0.287 | −0.059 | 4.5e−1 |
| Spectrin alpha, non-erythrocytic 1 | designed background | 43 | 3 | 9 | 0.664 | 0.721 | −0.057 | 1.3e−1 |
| Protein LG | designed background | 24 | 4 | 6 | 0.756 | 0.806 | −0.050 | 1.9e−1 |

| System[1] | Kind | Cells[2] | Cells via alanine | Deposit pairs | Mean ρ | Null mean ρ | Shift[3] | p[4] |
|---|---:|---:|---:|---:|---:|---:|---:|---:|
| SARS-CoV-2 receptor binding domain | natural variant | 31 | 3 | 4 | 0.363 | 0.413 | −0.050 | 3.1e−1 |
| SPTAN1 | designed background | 238 | 27 | 55 | 0.751 | 0.797 | −0.046 | 8.8e−4 |
| U14Ub1 | designed background | 50 | 4 | 8 | 0.556 | 0.597 | −0.041 | 3.4e−1 |
| ZFHX4 Homeobox domain | natural variant | 18 | 1 | 4 | 0.411 | 0.438 | −0.027 | 6.0e−1 |
| ZFHX3 Homeobox domain | natural variant | 19 | 1 | 4 | 0.439 | 0.462 | −0.024 | 6.0e−1 |
| PHOX2A Homeobox domain | natural variant | 49 | 3 | 4 | 0.404 | 0.420 | −0.016 | 6.0e−1 |
| UBC | designed background | 123 | 27 | 39 | 0.493 | 0.508 | −0.015 | 5.9e−1 |
| ALX3 Homeobox domain | natural variant | 46 | 5 | 5 | 0.392 | 0.404 | −0.012 | 6.2e−1 |
| FYN SH3 domain | designed background | 16 | 6 | 11 | 0.890 | 0.888 | +0.003 | 8.0e−1 |
| CHD5 Zinc finger, PHD-finger domain | natural variant | 9 | 0 | 3 | 0.301 | 0.298 | +0.003 | 8.0e−1 |
| ZFHX2 Homeobox domain | natural variant | 15 | 2 | 2 | 0.392 | 0.380 | +0.012 | 8.5e−1 |
| spg | designed background | 873 | 87 | 137 | 0.695 | 0.679 | +0.017 | 1.0e+0 |
| PAX7 Homeobox domain | natural variant | 53 | 2 | 4 | 0.422 | 0.402 | +0.020 | 9.7e−1 |
| ARX Homeobox domain | natural variant | 53 | 5 | 5 | 0.404 | 0.380 | +0.024 | 9.7e−1 |
| spa | designed background | 201 | 5 | 25 | 0.613 | 0.585 | +0.027 | 1.0e+0 |
| SARS-CoV-2 Omicron BA.1 receptor binding domain | natural variant | 21 | 1 | 4 | 0.486 | 0.458 | +0.028 | 9.7e−1 |
| ALX4 Homeobox domain | natural variant | 55 | 8 | 6 | 0.460 | 0.422 | +0.039 | 1.0e+0 |
| ZHER2 Affibody | designed background | 177 | 0 | 18 | 0.598 | 0.555 | +0.043 | 1.0e+0 |
| ALX1 Homeobox domain | natural variant | 45 | 7 | 5 | 0.499 | 0.456 | +0.043 | 1.0e+0 |
| entD | designed background | 8 | 3 | 6 | 0.839 | 0.788 | +0.051 | 9.7e−1 |

| System[1] | Kind | Cells [2] | Cells via alanine | Deposit pairs | Mean ρ | Null mean ρ | Shift[3] | p[4] |
|---|---|---|---|---|---|---|---|---|
| AR Zinc finger, nuclear hormone receptor-type domain | natural variant | 12 | 1 | 1 | 0.384 | 0.327 | +0.056 | 9.7e−1 |
| NR3C2 Zinc finger, nuclear hormone receptor-type domain | natural variant | 12 | 2 | 1 | 0.384 | 0.327 | +0.057 | 9.7e−1 |
| rplI | designed background | 48 | 0 | 15 | 0.923 | 0.848 | +0.075 | 1.0e+0 |
| Ice-structuring protein SP2 | designed background | 31 | 2 | 6 | 0.713 | 0.626 | +0.087 | 1.0e+0 |
| DMBX1 Homeobox domain | natural variant | 14 | 1 | 1 | 0.435 | 0.336 | +0.099 | 1.0e+0 |
| POU3F4 Homeobox domain | natural variant | 9 | 2 | 2 | 0.714 | 0.446 | +0.269 | 1.0e+0 |

[1]A system is the set of deposit pairs sharing a base target name, which groups engineered variants of one protein with its wild type and is not a clade, a family or a fold. Rows are ordered by the shift, so the systems composing worst against their own null come first.

[2]Collapsed onto one cell per distinct pair of target sequences and position. A system enters this table when it carries at least the number of cells the analysis plan requires.

[3]The mean rank correlation of the system minus the mean of its permutation null, so a negative value is a system composing worse than its own null.

[4]Permutation p, adjusted across the systems of this table by the Benjamini-Hochberg procedure.
